# Long-lasting astrocyte remodeling in Dravet Syndrome *Scn1a*^+/–^ mouse model

**DOI:** 10.64898/2026.01.06.697745

**Authors:** Athénaïs Genin, Alicia Janvier, Tristan Moujellil-Legagneur, Marine Blaquière, Alexis Chaussy, Romane Privé, Fabrice Duprat, Massimo Mantegazza, Etienne Audinat, Nicola Marchi, Noémie Cresto

## Abstract

**Background:** Dravet syndrome (DS) is a prototypical developmental and epileptic encephalopathy caused by *SCN1A* gene mutations leading to NaV1.1 loss of function. The latter causes early-onset drug-resistant seizures and enduring cognitive and behavioral deficits. In this pathological context, the implication of astrocytes remains insufficiently explored.

**Methods:** Using a heterozygous *Scn1a* knock-out (*Scn1a*^⁺/⁻^) mouse model that recapitulates the DS-human phenotype, we examine astrocyte remodeling at landmark disease stages, as defined by video-EEG and behavioral read-outs.

**Results:** From initial disease aggravation (PN20-35) to long-term stabilization (up to PN90), *Scn1a*^⁺/⁻^ mice showed increased hippocampal and cortical GFAP transcript and protein levels, compared to age-matched control littermates and to an earlier presymptomatic (<PN20) time point. During the aggravation phase in *Scn1a*^⁺/⁻^ mice, astrocyte branching, revealed by GFAP histological analysis and by intracellular delivery of Alexa Fluor 488 in hippocampal slices was increased but not sustained long-term. These disease-stage-dependent astrocyte modifications were not associated with macroscopic hippocampal sclerosis or cortical atrophy. To further study astrocyte remodeling during disease progression, we used biocytin diffusion following single-astrocyte loading to reveal an expanded astrocyte–astrocyte network in *Scn1a*^⁺/⁻^ mice long-term, along with increased Cx30 and Cx43 protein levels. An ethidium bromide uptake assay indicated impaired astrocytic hemichannel function in *Scn1a*^⁺/⁻^ mice long-term. Regionally, these long-term cellular and network astrocyte modifications coincided with augmented post-tetanic synaptic potentiation.

**Discussion:** In DS, astrocytes undergo a long-lasting network remodeling. We discuss how this astrocyte remodeling may be related to seizures as well as synaptic and cognitive deficits.

## Introduction

Developmental and epileptic encephalopathies (DEE) are severe early-onset disorders characterized by drug-resistant seizures and developmental delay ^1,2^. Dravet syndrome (DS), a prototypical DEE, is caused in most cases by *de novo* mutations in the *SCN1A* gene, which encodes the voltage-gated sodium channel Naᵥ1.1 ^3,4^. DS exemplifies how monogenic channelopathies can primarily drive epileptic activity and neurodevelopmental dysfunctions. It begins in the first year of life with severe, drugresistant seizures, leading to developmental delay and a high mortality rate, in particular because of status epilepticus and SUDEP. Bilateral or generalized tonic– clonic seizures persist into adulthood, when neurological deficits progress and plateau ^5–7^. Mouse models have been instrumental in showing that Naᵥ1.1 loss-of-function results in dysfunctional activity of inhibitory interneurons, leading to network hyperexcitability ^8–11^.

Here, we examine how genetically-driven seizures accompanied by progressive behavioral deficits in DS trigger, or are associated with, secondary responses: cellular modifications extending beyond neurons, with a particular focus on astrocytes ^12,13^. In some forms of epilepsy, such as temporal lobe and focal cortical seizures, astrocytes undergo a reactive transformation, which disrupts their homeostatic functions. These alterations can promote or sustain neuronal hyperexcitability, contributing to disease progression ^14^. However, in the specific context of DS, astrocyte reactivity and functional alterations have been only partially characterized, neither in the long-term nor at the functional level ^15^. Here, we used a heterozygous *Scn1a* knock-out (*Scn1a*^⁺/⁻^) mouse model haploinsufficient for NaV1.1 expression that recapitulates the temporal emergence and progression of seizures and cognitive deficits observed in DS patients^8^. Video-electroencephalography (V-EEG) and behavioral analyses were used to verify disease progression in this model. In this experimental framework, we studied glial cell reactivity, focusing on astrocyte remodeling, and obtained long-term histological, molecular, and network readouts. To confirm neuronal dysfunction overlapping with astrocyte remodeling ^16^, we acquired regional measures of synaptic activity using hippocampal slices. Our findings reveal that genetically driven seizures in DS induce long-lasting astrocytic remodeling, associated with persistent alterations in glial network properties and synaptic plasticity, thereby providing new insight into the cellular mechanisms underlying DS pathophysiology.

## Methods

### Animal housing

Heterozygous conditional global *Scn1a* knock-out (*Scn1a*^⁺/⁻^) mice were generated by crossing the *Scn1a* exon-25 floxed mouse line (*Scn1a^lox/lox^*) with mice expressing Cre recombinase under the Meox2 promoter (*Meox2-Cre^+^*) ^17–19^, both in the C57Bl/6J genetic background. *Scn1a^lox/+^:Meox-Cre****^−^*** mice were used as littermate controls. Mice were kept in standard mouse cages, under a 12h light/dark cycle, and allowed free access to food and water. Experiments were performed in accordance with national and European legislation, approved by ethical committees, supervised by the IGF local Animal Welfare Unit (A34-172–41) and by the French ministry (MESRI, APAFIS #50874-2024082809527395 v8). Genotyping was performed as in ^17^. Three clinically relevant DS progression stages are described in the *Scn1a*^⁺/⁻^ mouse model: 1) pre-symptomatic, before the appearance of seizures (<PN20); 2) an aggravation period with spontaneous seizures and related mortality as well as cognitive/behavioral defects (PN20–PN35); and 3) seizure stabilization with permanent cognitive and behavioral impairments (>PN35, long-term) (Suppl. Fig.1A). All examinations were performed at PN19, between PN27-PN32, and between PN85–PN95.

### Surgery, video-electroencephalography recordings, and analysis

At PN40-45, mice were subcutaneously injected with buprenorphine (0.05 mg/kg) 30 minutes before the procedure, followed by isoflurane anesthesia (induced with 5% isoflurane for 5 minutes and maintained at 2% in a mixture of 70% air and 30% O_2_). Body temperature was maintained at 37°C with a heating pad, and anesthesia depth was continuously monitored throughout the procedure. The scalp was shaved, prepared by cutaneous antisepsis using povidone–iodine, and locally anesthetized with a subcutaneous injection of lidocaine (4 mg/kg). The skin was then incised, the periosteum removed, and holes drilled for the placement of recording electrodes in the hippocampus, the frontal and parietal cortex (see Supp. Ext. Methods). The incision was closed with two 5/0 non-absorbable silk sutures (Ethicon), one placed anterior and the other posterior to the connector. Post-surgery, mice were monitored for well-being (weight loss, nest building, mobility, food/water intake, infection status, etc.). Enrichment, such as cotton nests and nest boxes, was always provided. Video-EEG was performed every two days in 12-hour recording sessions, alternating between night and day, from PN50 to approximately PN90 (see Supplemental Extended Methods for details).

### Immunofluorescence and quantifications

Mice were deeply anesthetized with ketamine/xylazine and perfused intracardially with saline solution. The brains were carefully dissected and post-fixed overnight with 4% PFA at 4°C. The next day, brains were cryoprotected (30% water, ethylene glycol, glycerol, and 10% 1M tris buffer) overnight in phosphate buffer saline solution (PBS) containing 30% sucrose. The brains were cut into 20 μm sections on a cryostat (Leica, Germany). Slices were stored in PBS with cryoprotectant at -20°C until use. For immunofluorescence, slices were blocked with PBS-Gelatin-Triton (PBS with 20% of horse serum and 0.25% of Triton) for 1h at room temperature, incubated overnight at 4°C with primary antibodies in the blocking solution, washed three times with PBS, incubated 2h with secondary antibodies in the blocking solution and washed three times with PBS before mounting with a Mounting Medium containing DAPI. Glial fibrillary acidic protein (GFAP) and ionized calcium-binding adaptor molecule 1 (IBA1) antibodies were used to stain astrocytes and microglia, respectively (see Supplemental Table 1). The secondary antibodies used were anti-chicken IgG conjugated to Alexa Fluor 488 and anti-rabbit IgG conjugated to Cy3. For morphological analysis of microglia and astrocytes, z-stack images were acquired using an automated imaging system (Axioscan, Zeiss, Plateforme Montpellier Ressources Imagerie) at 20X. The FIJI ′Sholl analysis’ plugin was then used to assess cell branching by measuring the number of intersections between GFAP or IBA1 staining and concentric circles spaced by 5 μm and centered on the cell nucleus. Astrogliosis and microglial reactivity were quantified by measuring GFAP and IBA1 immunoreactivity in the whole hippocampus using FIJI mean fluorescence intensity (MFI) measure. For microglial soma size quantification, a threshold was applied to select only microglial somas, followed by particle analysis in FIJI. Z-stack images of DAPI-stained sections (total thickness: 10 µm, comprising 17 optical slices of 0.60 µm each) were acquired to measure hippocampal and cortical layers. Regional maps were generated using an automated imaging system (AxioScan, Zeiss; 20× montage; Montpellier Ressources Imagerie). Four regions of interest (ROIs) were defined for analysis: hippocampal CA1, CA3, dentate gyrus (DG), and the somatosensory cortex. All analyses were conducted blind to the experimental group. The width of pyramidal cell layers in CA1 and CA3, the granule cell layer in the DG, and cortical thickness spanning layers I–VI (DAPI-positive nuclei) were quantified using FIJI. For each mouse, three sections were analyzed, and three measurements were obtained per section in CA1, CA3, and DG. See Supplementary Table 1 for all chemicals.

### Ex-vivo electrophysiological recordings and analysis

After rapid cervical dislocation, hippocampi were isolated and sectioned into 400 μm-thick slices in an ice-cold solution of artificial cerebrospinal fluid containing sucrose (aCSF sucrose composed in mM of 87 NaCl, 25 NaHCO_3_, 75 sucrose, 10 D-glucose, 2.5 KCl, 1 NaH_2_PO_4_, 7 MgCl_2_, 0.5 CaCl_2_ and oxygenated with carbogen (95% oxygen and 5% carbon dioxide) using a vibratome (VT1200S, Leica, Bannock-burn, IL, USA). The slices were stored at room temperature in a chamber containing the cutting solution for 15 min before being transferred to a second chamber containing normal aCSF (126 NaCl, 26 NaHCO_3_, 20 D-glucose, 2.5 KCl, 1.25 NaH_2_PO_4_, 1 Sodium pyruvate, 2 CaCl_2_ and 1 MgCl_2_ in mM and saturated with carbogen) heated to 34 °C, for at least 1 h before the first recording. Slices were transferred to an immersed chamber mounted on a BX51 Olympus microscope to record extracellular field potentials (fEPSP) (See Supplemental Extended Methods for details). To assess the detailed 3D morphology of astrocytes and the extent of their gap junctions coupling, CA1 astrocytes were whole-cell recorded with pipettes (5–6 MΩ) filled with a solution containing (in mm) 129 K-gluconate, 10 HEPES, 2 ATP-Mg^2+^, 10 EGTA and either 0.1 Alexa Fluor 488 (dextran, 3000 MW) or biocytin (20 mg/ml), adjusted to pH 7.3 with KOH. The dyes were passively loaded for 10 min during whole-cell recordings at -80 mV. The slices were then fixed in 4% paraformaldehyde before being imaged with a Leica SP8 confocal microscope (see Supplemental Extended Methods for details on electrophysiological recordings, image acquisition, and analysis).

### Ethidium bromide uptake assay

Slices were incubated in normal aCSF, in bursting solution (aCSF containing 0 mM Mg^2+^, 6 mM K^+^, and 100 µM picrotoxin), or in normal aCSF with Carbenoxolone (CBX, applied 30 min before and during EtBr incubation) containing EtBr. EtBr is a sensitive fluorescent tracer permeable to hemichannels that intercalates into DNA, thus becoming sequestered in the nucleus once absorbed by cells. After incubation, slices were rinsed for 10 minutes in appropriate aCSF, fixed for 12 hours in 4% paraformaldehyde in PBS (Ready-to-Use Fixative Solution, Biotium 22023), and immunostained for GFAP using a secondary anti-chicken IgG antibody conjugated to Alexa Fluor 488. Sections were then mounted with a DAPI-containing mounting medium. Image stacks were acquired with a Leica SP8 confocal microscope using a 40× objective at 1024×1024 resolution in sequential mode with excitation lasers set at 488 nm (GFAP), 561 nm (EtBr), and 350 nm (DAPI). Imaging was performed from the slice surface to the point of fluorescence extinction. GFAP-positive cells were analyzed to quantify astrocyte dye uptake. EtBr MFI was then measured using IMARIS software.

Detailed methods for qPCR, dye loading, electrophysiology, behavioral testing, and Western blotting are provided in the Supplemental Extended Methods.

### Statistics

Statistical analyses were performed using GraphPad Prism 10.2.2. Data normality was assessed with the Shapiro–Wilk test. Statistical tests are detailed in the figure legends. Two-way ANOVA or Kruskal–Wallis tests were used for normally or non-normally distributed data, respectively, with Bonferroni or Dunn’s post hoc corrections. Unpaired t-tests or Mann–Whitney tests were applied at individual time points. Cumulative distributions were compared using the Kolmogorov–Smirnov test. Data are presented as mean ± SEM. Survival was analyzed using Kaplan–Meier curves and log-rank (Mantel–Cox) tests. Correlations were computed in Python and reported as Pearson’s r with associated p-values.

## Results

### Neurophysiological hallmarks of experimental DS progression without overt tissue lesions

DS aggravation (tracked between PN20-PN30; Suppl.Fig.1A-B) was consistent with existing reports ^8,18,19^. Video-EEG monitoring during long-term stabilization (PN50 to PN90), conducted on a subset of mice (n=7 *Scn1a*^⁺/⁻^, 36 hours/week), captured generalized seizures (GS; Racine stage 4-5; Suppl. Video 1; Fig.1A and Suppl.Fig.1C) characterized by cortical-hippocampal discharges, with repetitions within minutes or hours depending on the individual mouse. These long-term epileptic patterns were associated with motor and cognitive deficits, as examined in all mice (PN85-90; Fig.1B1-B4). In the open field test, *Scn1a*^⁺/⁻^ mice exhibited hyperactivity compared with littermate controls, as evidenced by increased total distance travelled (Fig.1B1) that persists up to 6 months of age (Suppl.Fig.7C). In the Y-maze test, *Scn1a*^⁺/⁻^ mice showed a reduced percentage of spontaneous alternation, indicating impaired spatial working memory (Fig.1B2). In the three-chamber social interaction test, *Scn1a*^⁺/⁻^ mice spent similar amounts of time exploring the object and the mouse (Fig.1B3). Decreased marble burying corroborates the presence of autism-like traits (Fig.1B4).

**Figure 1.**
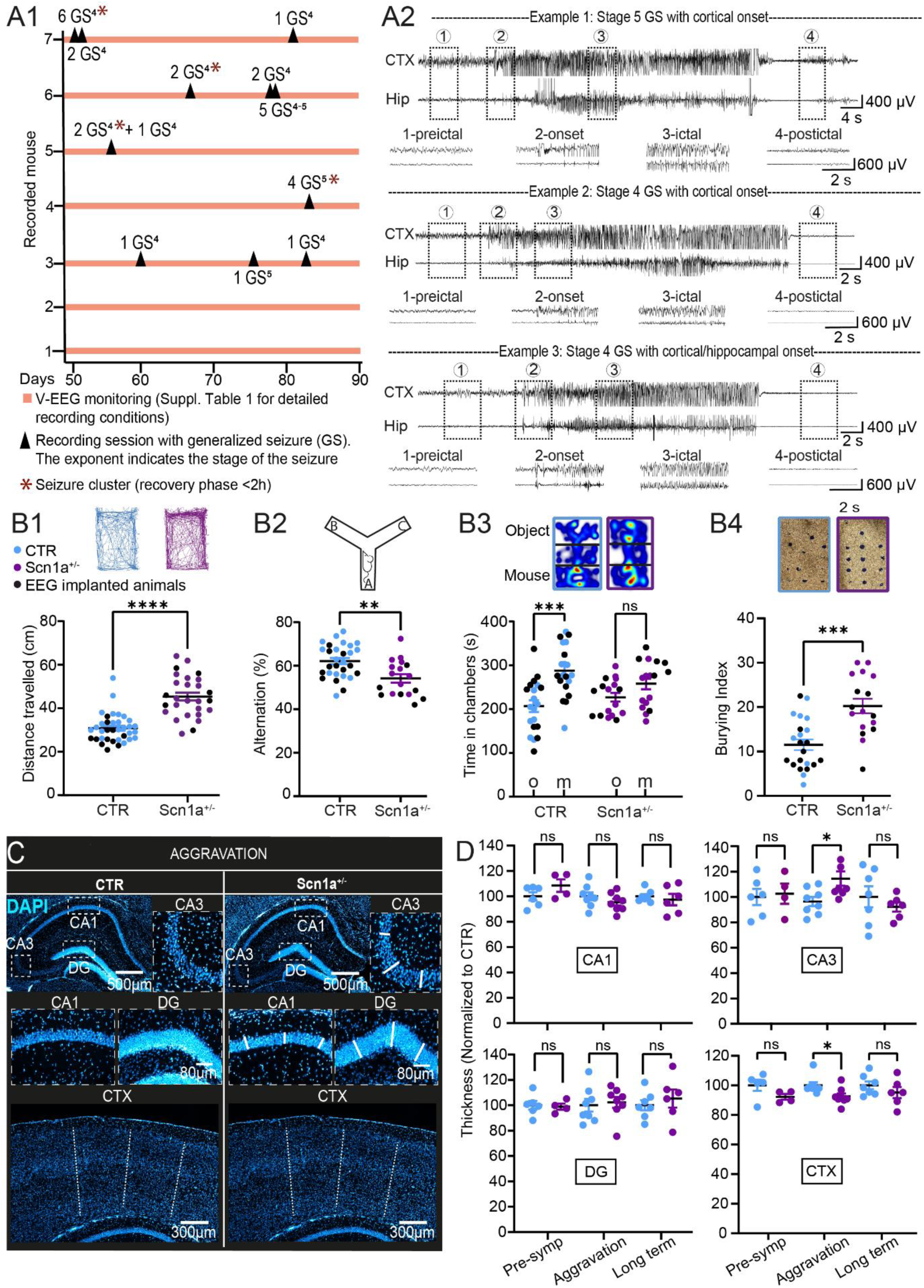
Spontaneous seizures and behavioral deficits in *Scn1a*^⁺/⁻^ mice do not provoke long-term hippocampal or cortical damage. **(A1)** *Scn1a*^⁺/⁻^ mice were individually monitored. For each animal, the initial Racine stage is indicated. Recordings were performed over 12 hours every two days, alternating between the light and dark phases, from PN50 to approximately PN90. **(A2)** Representative EEG traces and corresponding Racine stages. Examples include: a Stage 5 seizure with cortical onset (top; see zoom in panel 2), a Stage 4 seizure with cortical onset (middle; see zoom in panel 2), and a Stage 4 seizure with combined cortical/hippocampal onset (bottom; see zoom in panel 2). Insets (1), (3), and (4) illustrate pre-ictal, ictal, and postictal patterns, respectively. **(B)** Behavioral testing was performed long-term in both EEG-implanted and non-implanted mice (black dots). **(B1)** Locomotion was assessed in the open field test. Mann-Whitney test, CTR N=36; *Scn1a*^⁺/⁻^ N=27; p < 0.0001, **(B2)** Short-term memory was evaluated using the Y-maze in the alternation paradigm. Unpaired t test, CTR N=29; *Scn1a*^⁺/⁻^ N=18; p=0.0015 (t=3.377, df=45), **(B3)** Sociability was examined in the three-chamber test (object vs. mouse), Two-way ANOVA, CTR N=20; *Scn1a*^⁺/⁻^ N=17; CTR: p=0.0014; *Scn1a*^⁺/⁻^: p=0.3951, and **(B4)** marble burying behavior was also quantified, Unpaired t test, CTR N=22; *Scn1a*^⁺/⁻^ N=17; p=0.0001 (t=4.358, df=37). **(C)** Representative hippocampal DAPI-stained sections showing the absence of tissue sclerosis in *Scn1a*^⁺/⁻^ mice across stages. Examples of CA1, DG, and CA3 regions are shown (white lines indicate the measurement axes used for quantification). **(D)** Quantification of CA1, CA3, and DG thickness reveals no major pathological granule or pyramidal cell dispersion. Cortical thickness measurements across conditions are also shown. Unpaired T-test or Mann-Whitney tests were performed at each time point: Pre-symptomatic: CA1: CTR N=6; *Scn1a*^⁺/⁻^ N=4; p=0.1686, t=1.513, df=8; CA3: CTR N=6; *Scn1a*^⁺/⁻^ N=4; p=0.8067, t=0.2530, df=8; DG: CTR N=6; *Scn1a*^⁺/⁻^ N=4; p=0.8167, t=0.2396, df=8. CTX: Mann-Whitney test, CTR N=6; *Scn1a*^⁺/⁻^ N=6; p=0.1905. Aggravation: CA1: CTR N=6; *Scn1a*^⁺/⁻^ N=6; p=0.2675, t=1.155, df=14; CA3: CTR N=6; *Scn1a*^⁺/⁻^ N=4; p=0.0175, t=2.693, df=14; DG: CTR N=6; *Scn1a*^⁺/⁻^ N=4; p=0.7384, t=0.3407, df=14. CTX: Mann-Whitney test, CTR N=6; *Scn1a*^⁺/⁻^ N=6; p=0.0104. Long term: CA1: CTR N=7; *Scn1a*^⁺/⁻^ N=6; p=0.5853, t=0.5621, df=11; CA3: CTR N=6; *Scn1a*^⁺/⁻^ N=4; p=0.4284, t=0.8223, df=11; DG: CTR N=6; *Scn1a*^⁺/⁻^ N=4; p=0,5156, t=0.6718, df=11. CTX: Mann-Whitney test, CTR N=6; *Scn1a*^⁺/⁻^ N=6; p=0.3660. For all datasets, each dot represents one mouse. Statistical significance: ***p < 0.001; ****p < 0.0001.

We examined whether these neuropathophysiological trajectories are associated with brain morphological changes (Fig.1C-D). Quantification of DAPI maps ruled out hippocampal sclerosis at all time points examined (pre-symptomatic, disease aggravation, and stabilization) in *Scn1a*^⁺/⁻^ mice. Pyramidal and granular cell layer thickness was generally unchanged except for a modest increase in the CA3 cell dispersion at disease aggravation. Parietal cortex thickness was unmodified compared with age-matched littermate controls (Fig.1D). These results indicate that long-term spontaneous seizures and behavioral deficits do not provoke, or are not associated with, macroscopic structural modifications.

### Patterns of glial cell reactivity during experimental DS progression

In this pathophysiological context, we studied glial cell involvement initially by analyzing GFAP and IBA1 levels over time and across brain regions. Hippocampal (Fig.2A-D) and cortical (Suppl.Fig.2) GFAP transcript and immunoreactivity were increased at disease aggravation and long-term stabilization. No differences were found at an earlier, pre-symptomatic, time point. Astrocyte density (number/mm^2^) increased over time (Fig.2E), consistent with ^15^. Furthermore, Sholl analyses showed increased astrocytic branching complexity at disease aggravation in *Scn1a*^⁺/⁻^ mice, suggestive of reactive astrocytes at this disease stage ^20^ (Fig.2F). Hippocampal and cortical IBA1 levels and microglial soma size were increased specifically at disease aggravation, with no significant changes enduring in the long-term compared to agematched littermate controls (Suppl.Fig.3-4). Consistent with this mild reactive gliosis, we report that pro-inflammatory cytokine transcript levels (IL1β, IL6, TNFα, and CCL2) remained unchanged, except for a long-term increase in IL-6 (Suppl.Fig.5).

**Figure 2.**
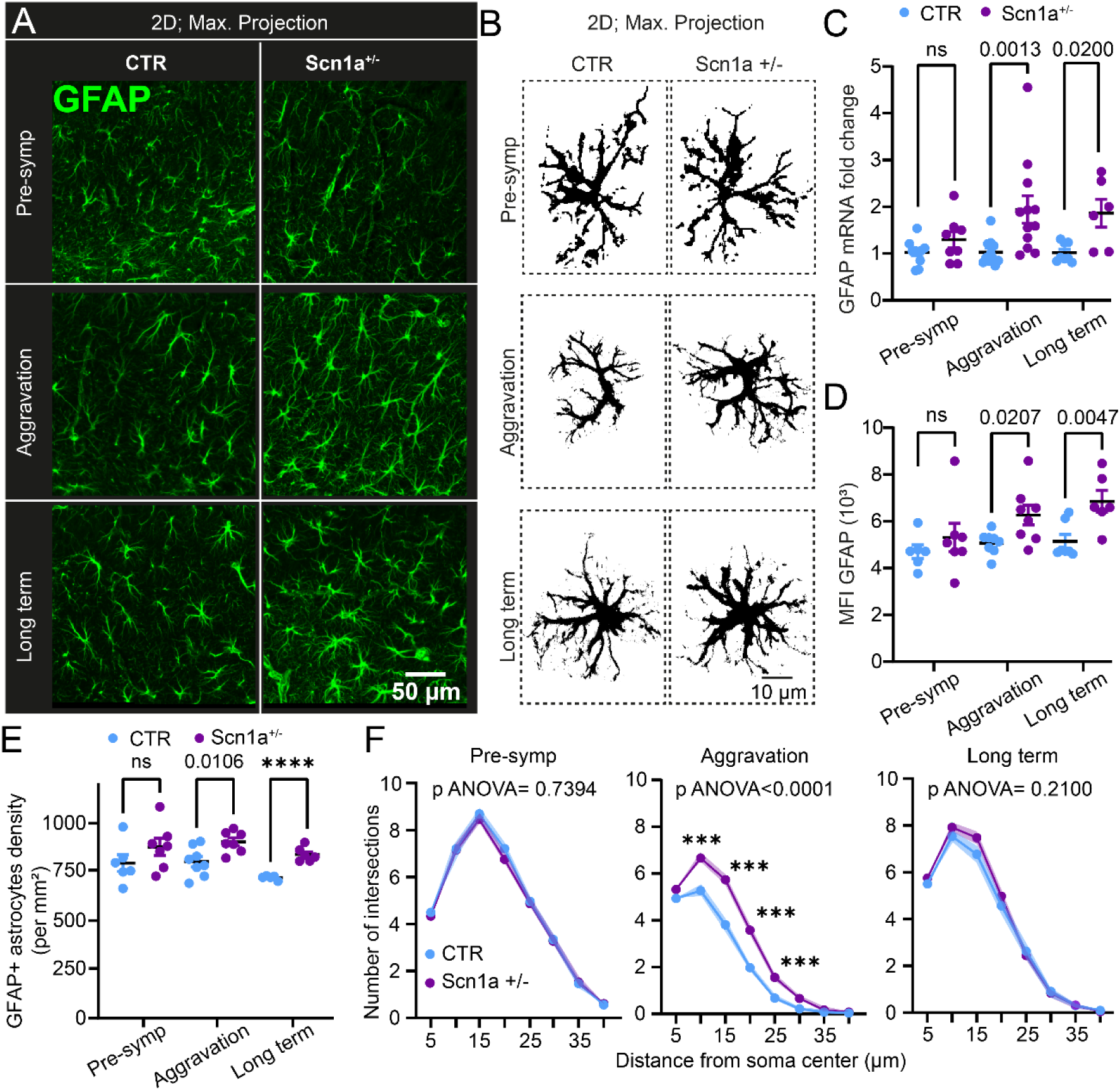
Long-term increase in hippocampal GFAP expression and astrocyte morphological remodeling in *Scn1a*^⁺/⁻^ mice. **(A-B)** Representative 40X GFAP images from the dorsal hippocampal CA1 region and magnification of individual astrocytes in CTR and *Scn1a*^⁺/⁻^ mice. Scale bar: 50 µm and 10 µm. **(C–D)** Quantification of GFAP mRNA levels (C) and GFAP mean fluorescence intensity (MFI) (D) at the pre-symptomatic, aggravation, and long-term phases. Mann–Whitney tests were performed at each time point: Pre-symptomatic: mRNA (CTR N=12; *Scn1a*^⁺/⁻^ N=8, p=0.2613), MFI (CTR n=6; *Scn1a*^⁺/⁻^ N=7, p=0.5338), aggravation: mRNA (CTR N=11; *Scn1a*^⁺/⁻^ N=12, p=0.0013), MFI (CTR N=8; *Scn1a*^⁺/⁻^ N=8, p=0.0207).Long term: mRNA (CTR N=8; *Scn1a*^⁺/⁻^ N=6, p=0.0200), MFI (CTR N=7; *Scn1a*^⁺/⁻^ N=6, p=0.0047). **(E)** Astrocyte density quantified from GFAP immunolabelling in the stratum radiatum. Unpaired t-tests were performed at each time point: Pre-symptomatic: CTR N=6; *Scn1a*^⁺/⁻^ N=7; p=0.2053, t=1.346, df=11. Aggravation: CTR N=8; *Scn1a*^⁺/⁻^ N=7; p=0.0106, t=2.980, df=13. Long term: CTR N=5; *Scn1a*^⁺/⁻^ N=6; p<0.0001, t=6.904, df=9. **(F)** Sholl analysis was performed on individual CA1 astrocytes at each age. Intersections were quantified using concentric circles spaced 5 µm apart from the soma. Two-way ANOVA was used at each time point: Pre-symptomatic: CTR 100 cells (N=5); *Scn1a*^⁺/⁻^ 100 cells (N=5); interaction p=0.7394, F(4.458, 882.8)=0.5218. aggravation: CTR 100 cells (N=5); *Scn1a*^⁺/⁻^ 100 cells (N=5); interaction p<0.0001, F(3.292, 782.5)=11.89; followed by Bonferroni’s post hoc test. Long term: CTR 100 cells (N=5); *Scn1a*^⁺/⁻^ 100 cells (N=5); interaction p=0.2100, F(7, 1175)=1.380. For all datasets, each dot represents one mouse. Statistical significance: ***p < 0.001; ****p < 0.0001.

Building on this initial examination and to further investigate astrocyte changes specifically during disease aggravation and in the long-term, we assessed three-dimensional branching morphology by intracellular loading of single astrocytes with the fluorescent tracer Alexa Fluor 488 in ex vivo hippocampal slices. Examples of the 3D astrocyte territory and architecture are shown in Fig.3A. Using this approach, we identified astrocytic morphological reactivity during disease aggravation in *Scn1a*^⁺/⁻^ mice (Fig. 3B1–B2), which was not maintained long-term (Fig.3C1–C2), based on Sholl analysis and total cell volume measurements (Fig.3B3–C3), relative to age-matched littermate controls. We report increased astrocyte soma volume over time (Fig.3B4-C4). Overall, these results portray specific morphological modifications of astrocytes during DS aggravation and in the long-term, with regional changes reflecting the generalized seizure phenotype and behavioral deficits that develop over time (see Fig.1).

**Figure 3.**
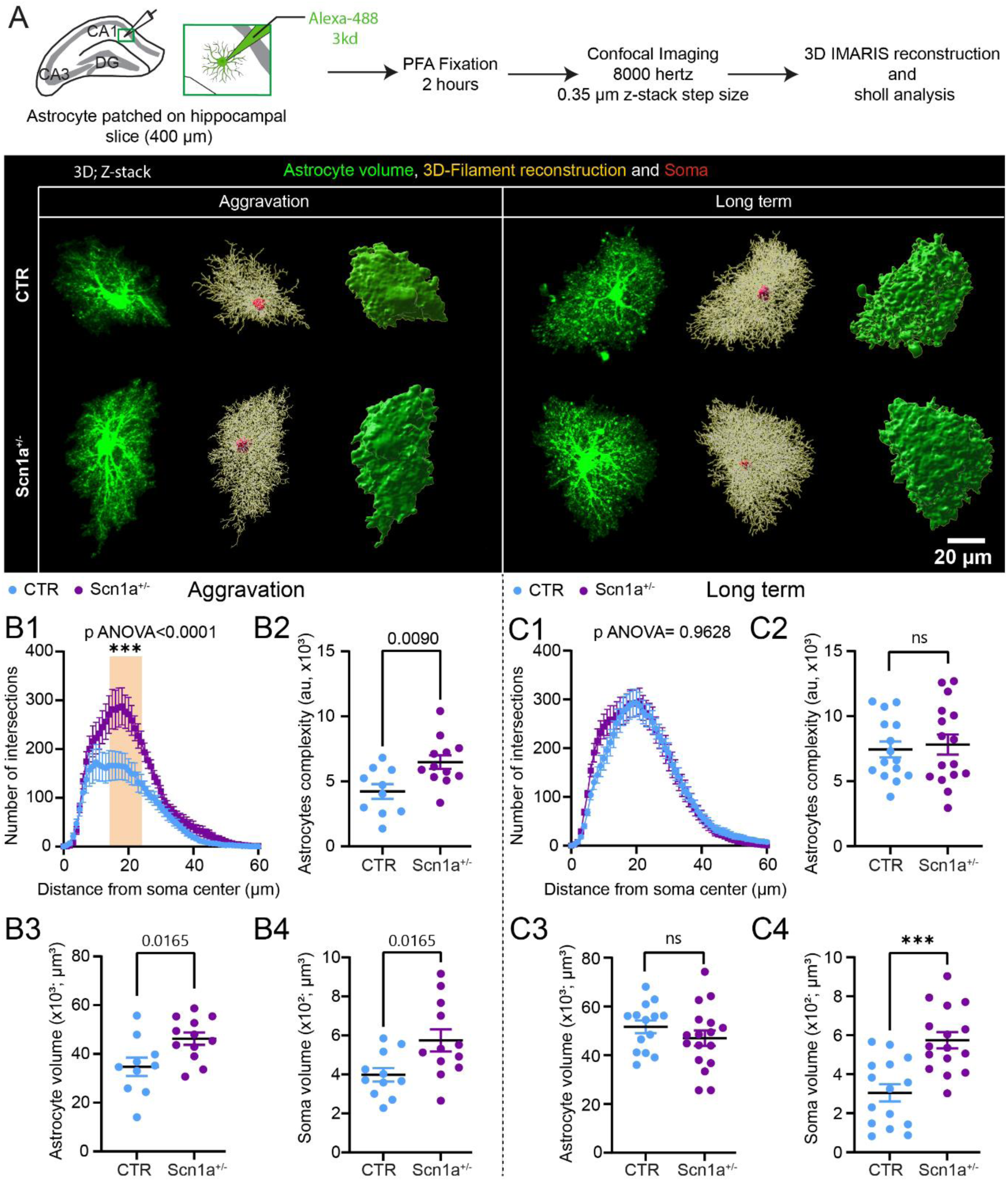
Three-dimensional astrocyte morphological changes in *Scn1a*^⁺/⁻^ mice. **(A)** Diagram showing Alexa-488 dye loading and spread throughout astrocytic processes, acquisition of high-resolution z-stacks, and subsequent 3D reconstruction. Representative 40× images of Alexa-488–loaded astrocytes in the CA1 region of CTR and *Scn1a*^⁺/⁻^ mice at aggravation and long-term phases. Scale bar: 20 µm. Astrocytes were reconstructed in 3D using IMARIS software, filaments and volumes are represented. **(B1-C1)** Sholl analysis of individual Alexa-488–loaded astrocytes at each age. Intersections were quantified using concentric circles spaced 1 µm apart from the soma in the 3D module of IMARIS. Two-way ANOVA was performed at each time point: Aggravation: CTR 10 cells (N=5); *Scn1a*^⁺/⁻^ 12 cells (N=6); interaction p<0.0001, F(60, 1202)=2.285; followed by Bonferroni’s post hoc test. Statistical differences were observed from 14 µm to 24 µm from the soma (14 µm: p=0.0234; 15 µm: p=0.0055; 16 µm: p=0.0010; 17 µm: p=0.0012; 18 µm: p=0.0006; 19 µm: p=0.0003; 20 µm: p=0.0016; 21 µm: p=0.0011; 22 µm: p=0.0030; 23 µm: p=0.0281; 24 µm: p=0.0257). Long term: CTR 15 cells (n=10); *Scn1a*^⁺/⁻^ 16 cells (n=9); interaction p=0.9628, F(60, 1037)=0.6943. **(B2-C2)** Astrocyte complexity was quantified as the area under the curve from Sholl analysis. Aggravation **(B2):** Unpaired t-test, p=0.0090, t=2.894, df=20. Long term **(C2):** Unpaired t-test, p=0.7048, t=0.3826, df=29. **(B3-C3)** 3D astrocytic volume. Aggravation **(B3):** Unpaired t-test, p=0.0165, t=2.618, df=20. Long term **(C3):** Unpaired t-test, p=0.2766, t=1.109, df=29. **(B4-C4)** 3D astrocytic soma volume. Aggravation **(B4):** Unpaired t-test, p=0.0165, t=2.607, df=21. Long term **(C4):** Unpaired t-test, p=0.0001, t=4.461, df=29. For all datasets, each dot represents one astrocyte. Statistical significance: ***p < 0.001.

### The astrocyte network is remodeled in experimental DS

Next, we examined whether astrocytic changes accompanying seizure aggravation and long-term disease in *Scn1a*^⁺/⁻^ mice were associated with remodeling of the astrocyte–astrocyte network. This query is relevant given astrocytes’ buffering capacity, which depends on intercellular communications ^21^. We quantified the 3D astrocytic network by intracellularly loading a single astrocyte with biocytin, a tracer capable of diffusing across inter-cellular gap-junctions in *ex vivo* hippocampal slices (Fig.4A). Using a python-based routine, we generated 3D astrocyte maps across experimental conditions (Fig.4B). We report a significantly increased number of astrocyte coupling in *Scn1a*^⁺/⁻^ mice compared to age-matched control littermates (aggravation phase: 47.56 +/- 2.940 and 65.5 +/- 3.888 for control and *Scn1a*^⁺/⁻^, respectively; long-term stabilization: 43.00 +/- 3.145 and 81.75 +/- 6.044 for control and *Scn1a*^⁺/⁻^, respectively; Fig.4C). Next, we calculated the 3D cellular envelope to examine the total volume occupied by the astrocytic network. We report a significantly increased envelope volume of astrocyte networks in *Scn1a*^⁺/⁻^ mice in the long-term, while no changes were detected in control littermates (Fig.4D). The number of astrocytes directly coupled to the patched astrocyte was approximately four (Fig.4E-F), with increased inter-astrocyte proximity in *Scn1a*^⁺/⁻^ mice (Fig.4G).

**Figure 4.**
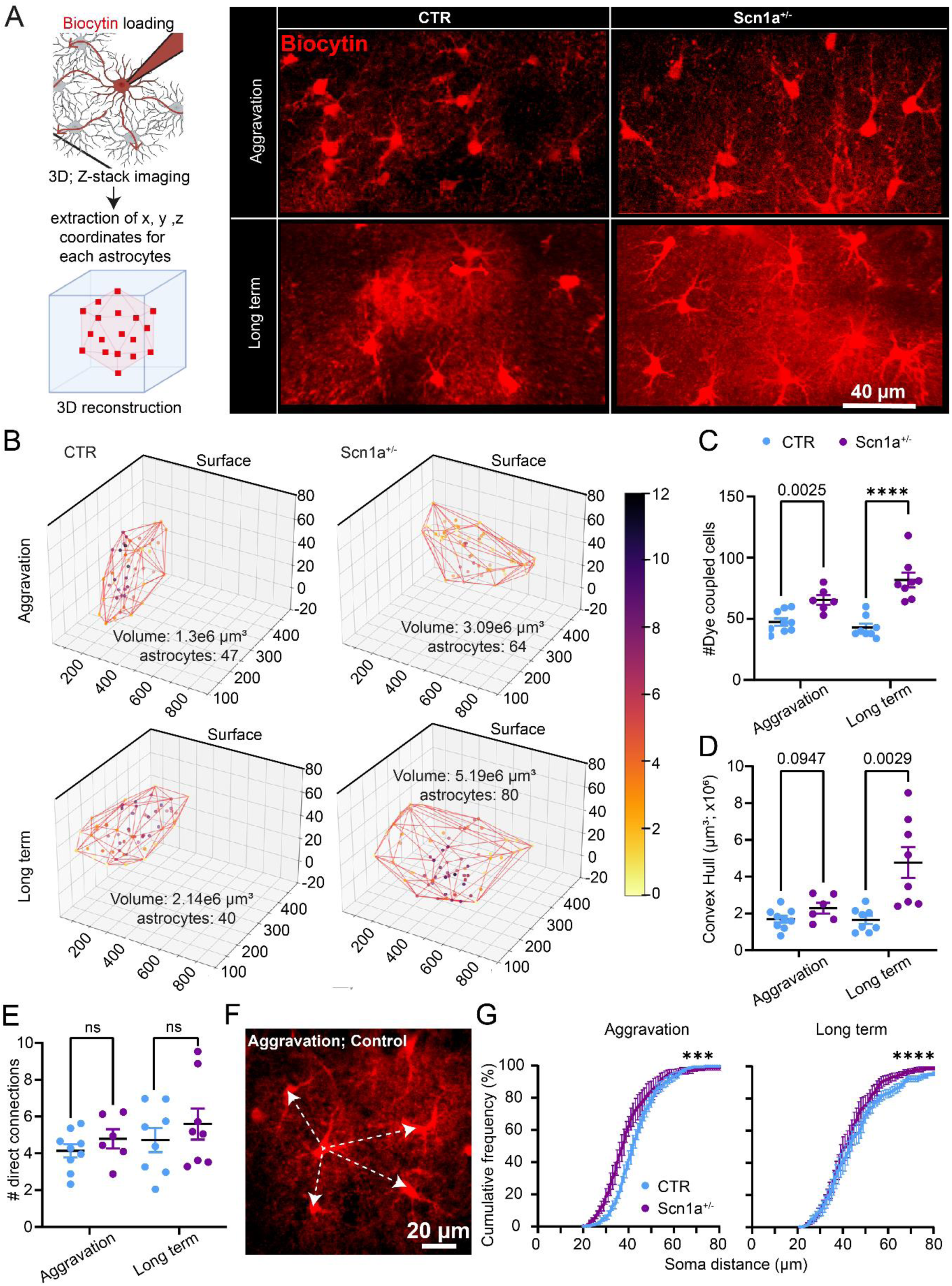
Increased astrocyte network connectivity during disease progression in *Scn1a*^⁺/⁻^ mice. **(A)** Diagram showing biocytin loading and spread throughout the CA1 astrocytic syncytium, acquisition of high-resolution z-stacks, and subsequent 3D reconstruction of the network based on centroid coordinates extracted for each astrocyte. Representative 40× images of biocytin-loaded astrocytes in the CA1 region of control (CTR) and *Scn1a*^⁺/⁻^ mice at aggravation and long-term stages. Scale bar: 40 µm. **(B)** Astrocyte networks were reconstructed in 3D using IMARIS software combined with a custom Python script. **(C–D)** Quantification of the number of coupled cells (C) and the convex hull volume (D). **(C)** Number of coupled cells: Aggravation: unpaired t-test, p = 0.0025, t = 3.743, df = 13; CTR n = 9 mice; *Scn1a*^⁺/⁻^ N = 6 mice. Long term: unpaired t-test, p < 0.0001, t = 5.687, df = 14. CTR N = 8 mice; *Scn1a*^⁺/⁻^ N = 8 mice. **(D)** Convex hull volume: Aggravation: unpaired t-test, p = 0.0947, t = 1.802, df = 13; CTR N = 9 mice; *Scn1a*^⁺/⁻^ N = 6 mice. Long term: unpaired t-test, p = 0.0029, t = 3.604, df = 14. CTR N = 8 mice; *Scn1a*^⁺/⁻^ N = 8 mice. **(E)** The number of directly connected cells was quantified using a Python script. Aggravation : unpaired t-test, p = 0.3070, t = 1.063, df = 13 ; CTR N = 9 mice; *Scn1a*^⁺/⁻^ N = 6 mice. Long term: unpaired t-test, p = 0.4270, t = 0.8182, df = 14. CTR N = 8 mice; *Scn1a*^⁺/⁻^ N = 8 mice. **(F-G)** Representative image of CTR hippocampal slice from the aggravation phase, and the distance between the four closest neighbours quantified and plotted as cumulative frequency distributions. Scale bar: 20 µm. Kolmogorov–Smirnov test: Aggravation, p = 0.0008; Long term, p < 0.0001.

The long-term reorganization of the astrocytic network prompted us to investigate hippocampal Cx30 and Cx43 expression, the primary proteins that form the gap junctions ^22^. Consistent with the long-term expansion of the astrocyte network, *Scn1a*^⁺/⁻^ mice show increased levels of Cx30 and Cx43 (Fig.5A-C). We examined the phosphorylated form of Cx43, which is known to regulate gap junction function. We report a non-significant trend toward an increase in *Scn1a*^⁺/⁻^ mice (Fig.5D). Within this framework, we confirmed GFAP increase in *Scn1a*^⁺/⁻^ mice in the long-term (Fig.5E) and up to 6 months of age (Suppl.Fig.7A-B). Next, we examined mouse-specific correlations between long-term behavioral deficits and Cx and GFAP levels. We report significant positive correlations between levels of Cx30, Cx43, and phosphorylated Cx43 in the hippocampus and distance traveled in the open field (Fig.5F). Cx30 levels positively correlated with deficits in marble burying. No significant correlation was observed between the tested proteins and performance in the Y-maze. These findings link astrocytic alterations and connexin expression to specific long-term behavioral outcomes in Scn1a⁺/⁻ mice, while indicating that short-term memory performance (Y maze) may be dissociable from the astrocytic dysfunctions assessed here.

**Figure 5.**
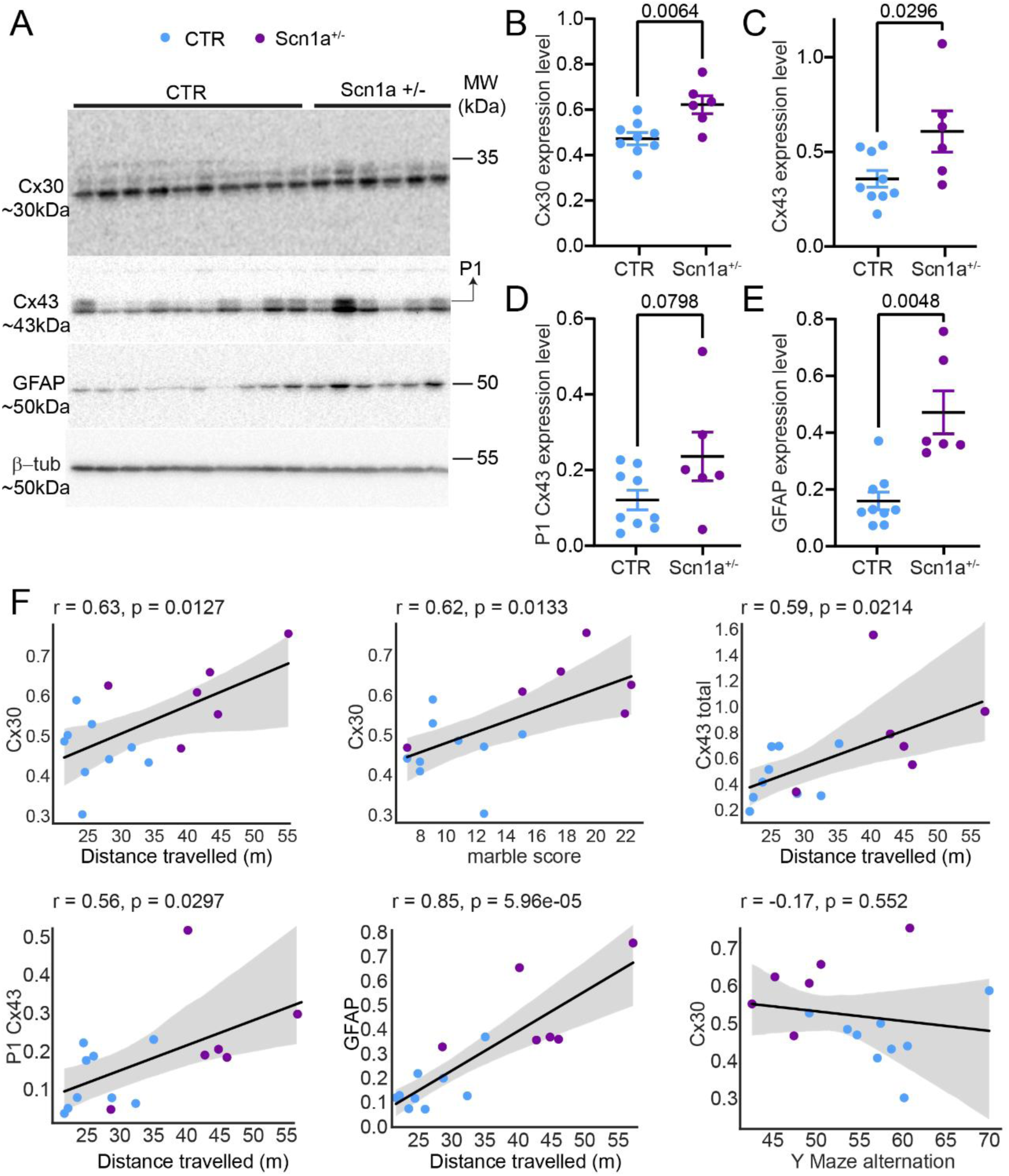
Increased expression of Cx30 and Cx43 long-term in *Scn1a*^⁺/⁻^ mice. **(A)** Western blot analysis of Cx30, Cx43, GFAP, and β-tubulin protein levels in CTR (N = 9) and *Scn1a*^⁺/⁻^ mice (N = 6) in the long-term phase. P1 bands correspond to phosphorylated forms of Cx43. **(A–E)** Quantification of Cx30, Cx43, phosphorylated (P1) form of Cx43, and GFAP (B; unpaired t-test, p = 0.0064, t = 3.242, df = 13. C; unpaired t-test, p = 0.0296, t = 2.443, df = 13. D; unpaired t-test, p = 0.0798, t = 1.900, df = 13. E; Mann-Whitney test, p = 0.0048), normalized to β-tubulin. **(F)** Correlation between Cx30 (r = 0.63, p = 0.0127), Cx43 (r = 0.59, p = 0.0214), phospho Cx43 (r = 0.56, p = 0.0297), GFAP (r = 0.85, p = 0.0000596) and distance travelled in the open field. Cx30 levels correlate with the marble-burying score (r = 0.62, p = 0.0133). Cx30 levels do not correlate with the Y-maze alternation (r = -0.24, p = 0.38).

### Astrocyte hemichannel and synaptic dysfunction in DS

Because connexins form hemichannels involved in gliotransmitter release and regulation of neuronal excitability ^23^, we assessed ethidium bromide (EtBr) cellular uptake, a functional biomarker. In hippocampal slices, under normal artificial CSF conditions, *Scn1a*^⁺/⁻^ mice and control littermates showed EtBr astrocytic uptake. Whereas bursting conditions (0 Mg²⁺ and 6 mM K⁺) expectedly increased EtBr uptake in control littermates, uptake in *Scn1a*^⁺/⁻^ mice was reduced to levels comparable to those produced by CBX, a hemichannel blocker (Fig.6A-C). These results suggest impaired hemichannel functionality during epileptiform discharges in hippocampal slices obtained from *Scn1a*^⁺/⁻^ mice. Finally, to reveal neuronal dysfunction concomitant with astrocyte remodeling, we evaluated synaptic activity in the CA1 hippocampal region where this remodeling occurred. In *Scn1a*^⁺/⁻^ and control mice (Fig.6D), upon Schaffer collateral stimulations, CA1 basal synaptic transmission and paired-pulse facilitation were comparable (Fig.6D-E). The initial response after tetanic stimulation was greater in *Scn1a*^⁺/⁻^ slices, while overall LTP was similar between genotypes (Fig.6F). To isolate post-tetanic potentiation (PTP), we blocked post-synaptic NMDA receptors with D-AP5 (50 µM). We observed a significantly enhanced PTP in *Scn1a*^⁺/⁻^ mice over the long-term (Fig.6G) and up to 6 months (Suppl.Fig.7D), suggesting increased glutamate release during tetanic stimulation. Finally, to test whether a larger readily releasable vesicle pool contributes to this effect, we applied 10 Hz stimulation. *Scn1a*^⁺/⁻^ slices displayed stronger initial facilitation and reduced subsequent depression (Fig.6H). These findings portray the simultaneous presence of regional astrocyte remodeling and altered synaptic transmission properties in the long-term in experimental DS.

**Figure 6.**
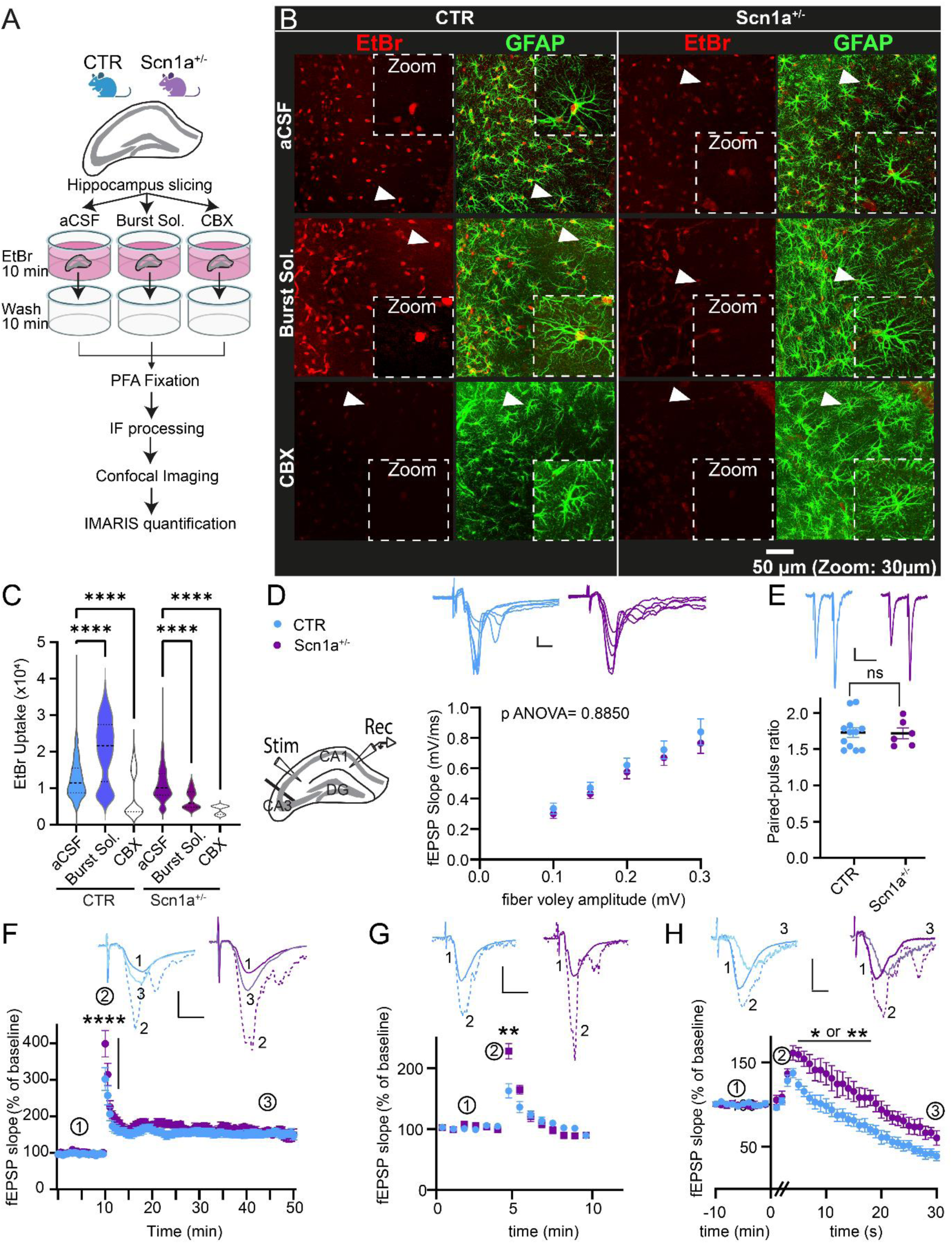
Long-term reduction of astrocytic hemichannel function and synaptic defects in *Scn1a*^⁺/⁻^ mice. **(A)** Hemichannel (HC) opening was assessed using the HC-permeable fluorescent tracer ethidium bromide (EtBr; 314 Da, 4 µM, 10 min) in acute hippocampal slices. Slices were incubated for 1h in either normal aCSF, bursting solution (0 Mg–6 K–P; 100 µM; applied 30 min before and throughout EtBr exposure), or aCSF containing carbenoxolone (CBX; 200 µM; applied 30 min before and during EtBr incubation). After labeling, slices were rinsed for 10 min in aCSF, fixed for 12 h in 4% paraformaldehyde, immunostained for GFAP, and mounted in ProLong™ Gold with DAPI. Scale bar: 50 µm. **(B)** Representative images of EtBr (red) and GFAP (green) in the CA1 region of control (CTR) and *Scn1a*^⁺/⁻^ mice in long-term phase under aCSF, bursting, and CBX conditions. Arrows indicate the astrocyte shown in the zoomed panels. **(C)** Dye uptake was quantified by measuring EtBr fluorescence intensity in individual astrocytes using IMARIS (CTR aCSF: n = 2242 astrocytes from 13 mice; CTR burst: n = 315 from 3 mice; CTR CBX: n = 282 from 3 mice; *Scn1a*^⁺/⁻^ aCSF: n = 999 from 10 mice; *Scn1a*^⁺/⁻^ burst: n = 120 from 2 mice; *Scn1a*^⁺/⁻^ CBX: n = 13 from 2 mice). Mann–Whitney test; ****p < 0.0001. **(D)** Schematic showing electrode placement in the stratum radiatum. Schaffer collaterals (SC) were stimulated and fEPSPs recorded in CA1. Input–output curves indicate unchanged basal excitatory synaptic transmission in *Scn1a*^⁺/⁻^ mice compared to CTR at PN90 (CTR: N = 11 mice; *Scn1a*^⁺/⁻^: N = 6 mice; two-way ANOVA, p = 0.8850). Scale bar: 0.2 mV; 10 ms. **(E)** Paired-pulse facilitation (PPF) was also unchanged in *Scn1a*^⁺/⁻^ mice (CTR: N = 10 mice; *Scn1a*^⁺/⁻^: N = 7 mice; Mann–Whitney test, p = 0.9192). Scale bar: 0.1 mV; 40 ms. **(F)** Long-term potentiation (LTP) at CA1 Schaffer collateral synapses was recorded in PN90 *Scn1a*^⁺/⁻^ mice (N = 7) and CTR littermates (N = 10). fEPSPs after the stimulation train (trace 3) were normalized to baseline (trace 1). Sample traces show averaged fEPSPs before (trace 1), immediately after tetanization (dashed line, trace 3), and 50 min post-tetanization (light color). Two-way ANOVA: genotype effect p < 0.0001; F(1,1515) = 75.30. **(G)** Post-tetanic potentiation (PTP) was increased in *Scn1a*^⁺/⁻^ mice (N = 7) compared to CTR (N = 8). Two-way ANOVA revealed a significant genotype × time interaction (p < 0.0001; F(3.233, 42.03) = 10.78). Bonferroni post hoc: p = 0.0075. Sample traces show averaged fEPSPs before and immediately after tetanization (dashed line). **(H)** The readily releasable vesicle pool was evaluated using a 10 Hz, 30 s stimulation protocol and normalized to baseline (5 min, trace 1). Scale bar: 0.4 mV; 5 ms. Two-way ANOVA revealed significant differences between groups from the 5th to the 18th second (except for two time points):5 s: p = 0.0270; 6 s: p = 0.0102; 7 s: p = 0.0044; 8 s: p = 0.0123; 9 s: p = 0.0069; 10 s: p = 0.0112; 11 s: p = 0.0167 / 0.0159; 12 s: p = 0.0194; 13 s: p = 0.0455; 16 s: p = 0.0151. Asterisks indicate significance (*p < 0.05; **p < 0.01; ***p < 0.001; ****p < 0.0001).

## Discussion

Our results show how astrocyte remodeling occurs over the long-term in a *Scn1a*^⁺/⁻^ mouse model of DS, driven by seizures induced by the genetic condition and correlating with cognitive deficits. Given their homeostatic functions, astrocytes may represent an entry point for a better understanding of the pathological mechanisms in DS, with a possible extension to other DEE. Together with the stable increase in GFAP levels, specifically during the DS aggravating stage, we report histological signs of astrocyte reactivity, likely a consequence of severe seizures at this initial and critical disease stage (Fig.2-3; Suppl.Fig.2-4), as also proposed in ^15,24^. Furthermore, existing studies report either no microglial alterations ^25^ or a mild transient activation during the early phase of the disease ^15,24^, findings that agree with our results (Suppl.Fig.3-4). In the long-term, during disease stabilization, we showed that only astrocytes were remodeled, as indicated by their extended network (Fig.4), while no microglial involvement was detected at least with the analyses we performed ^26,27^.

Importantly, the magnitude of reactive gliosis herein observed was lower than that typically reported in experimental models of temporal lobe epilepsy (TLE), consistent with the absence of overt hippocampal sclerosis ^28^. Thus, although the latter can drive glial activation in lesional epileptogenesis and in chronic focal seizures, it is absent in the DS model. Consistent with this non-lesional DS framework, pro-inflammatory cytokine levels did not change over time, except for IL-6 (Suppl.Fig.5). From these data, we infer that the observed long-term astrocyte remodeling does not imply a stable pro-inflammatory trajectory; instead, it could reflect a homeostatic adaptation to abnormal neuronal activity in the long-term.

### Astrocyte remodeling and connexin expression in DS

A significant finding of our work is the augmentation of the astrocyte network from DS aggravation to long-term (Fig.4). We also report increased long-term expression of Cx30 and Cx43 (Fig.5). The latter is consistent with previous data from adult human autopsy DS brains ^12^. Notably, absence epilepsy models show increased connexin expression and astrocytic coupling, potentially promoting network synchronization ^29,30^. TLE models display more heterogeneous outcomes, with reports of increased ^31–34^, unchanged ^35–37^, or decreased levels of Cx30 and Cx43 ^38,39^, as well as a study demonstrating subcellular redistribution of Cx43 and functional astrocytic uncoupling despite preserved protein levels ^40^. Another study reported a complete loss of functional gap junction coupling between astrocytes in the sclerotic hippocampus from drug-resistant TLE patients ^41^. Based on our results and existing evidence, we suggest that the trajectory of astrocyte network remodeling and Cx expression over time are contingent on lesion presence and seizure type, as in focal vs. generalized genetic seizures.

### Astrocytes and neurophysiological adaptations in DS: is there a link?

Long-term changes in the astrocytic network could be associated with cognitive deficits ^42^, as reported here in DS. Experimental manipulation of astrocytic connexins has provided insights into how network size and intercellular coupling shape neuronal function. Upregulation of Cx30 enlarges the astrocytic network, dampens spontaneous and evoked synaptic transmission, and impairs recognition memory ^43^. Additionally, viral-mediated manipulations of Cx43 demonstrated that fine-tuning connexin expression can modulate cognition and excitability ^44,45^.

In contrast with enhanced gap junction coupling and increased expression of Cx30 and Cx43 (Fig.4-5), we observed reduced hemichannel function in *Scn1a*^⁺/⁻^ mice (Fig.6). Hemichannel opening can contribute to enhancing neuronal excitability by releasing ATP and glutamate, and has been implicated in the generation of pilocarpine-induced acute seizures ^46–49^. Our results thus support the hypothesis of a shift toward adaptive gap-junctional coupling and limiting potentially harmful hemichannel opening in experimental DS ^46^.

Previous work showed that hippocampal overexpression of Cx30 reduced posttetanic potentiation ^43^. Here, we report that in *Scn1a*^⁺/⁻^ mice, increased Cx30 expression and astrocyte gap junction coupling are associated with an increased posttetanic potentiation in DS. This highlights the complex and context-dependent role of Cx30 in modulating synaptic plasticity. Importantly, we did not detect any change in the number of excitatory synapses, indicating that structural synapse addition is unlikely to account for the enhanced post-tetanic potentiation (Suppl.Fig.6).

### Study limitations and perspectives

Our study has limitations. First, we did not control for the timing of the last seizure when evaluating astrocytic and microglial histological activation. In the context of DS, it remains to be determined whether spontaneous generalized seizures, occurring over the long-term, are sufficient to elicit an acute, potentially transient glial activation pattern. Such activation could drive glial-omics alterations, leading to adaptive responses that may be either beneficial or detrimental during disease progression and contributing to the astrocytic network remodeling observed in this study. Second, we are aware that the data presented do not permit establishing a causal relationship or a direct functional interaction between increased astrocytic network remodeling, the long-term persistence of hippocampal synaptic deficits, and behavior or cognitive changes. One potential mechanism involves dysregulation of astrocytic glutamate homeostasis; however, this warrants further investigation. Finally, potential sex-specific differences were not systematically addressed. Our analyses were limited to selected brain regions and disease stages, and did not directly assess dynamic astrocyte functions.

### Conclusions

In summary, genetically driven seizures, in the absence of overt tissue damage, are sufficient to promote long-term, stable astrocyte remodeling, associated with behavioral deficits. In this framework, we propose that astrocytes participate in the progression of experimental DS, exhibiting phenotypes that evolve over time, from early morphological alterations to long-term network-level changes. The precise molecular players underlying these modifications and whether a disease-related direct glio–neuronal functional link exists remain to be fully elucidated.

## Supporting information

Supplemental video Stage 5

## Acknowledgments

This work was supported, as a whole or in part, by FFRE, Chamaillard, ANR-DSRemo (ANR-24-CE14-3636), ANR-EpiNeurAge (anr-22-ce14-0042), ANR-EpiCatcher (ANR-21-CE17-0031), ANR-CEST-Focus (ANR-22-CE17-0061), France 2030 investment plan as part of the Université Côte d’Azur’s Initiative of Excellence (ANR-15-IDEX-01), Université Côte d’Azur Strategic IdEx program PSI-Ion Channels. We thank Emilie Bonnet (Institute of Molecular and Cellular Pharmacology, University Cote d’Azur, CNRS UMR 7275 Inserm U1323) for her skillful technical support on colony management/genotyping.

## Conflict of Interest/Ethical Publication Statement

None of the authors has any conflict of interest to disclose.

## Ethical publication

We confirm that we have read the Journal’s position on issues involved in ethical publication and affirm that this report is consistent with those guidelines.

## Authors’ contributions

Author Contributions. Conceptualization: A.G., E.A., N.M., N.C.; Formal analysis: A.G., A.J., T.M.L., A.C., R.P., M.B, N.C.; Funding acquisition: M.M., E.A., N.M., and N.C.; Supervision: N.C.; Project administration: N.C.; Writing original draft: N.C., A.G., N.M.; Writing – review & editing: all authors.

---

**Supplemental Figure 1.**
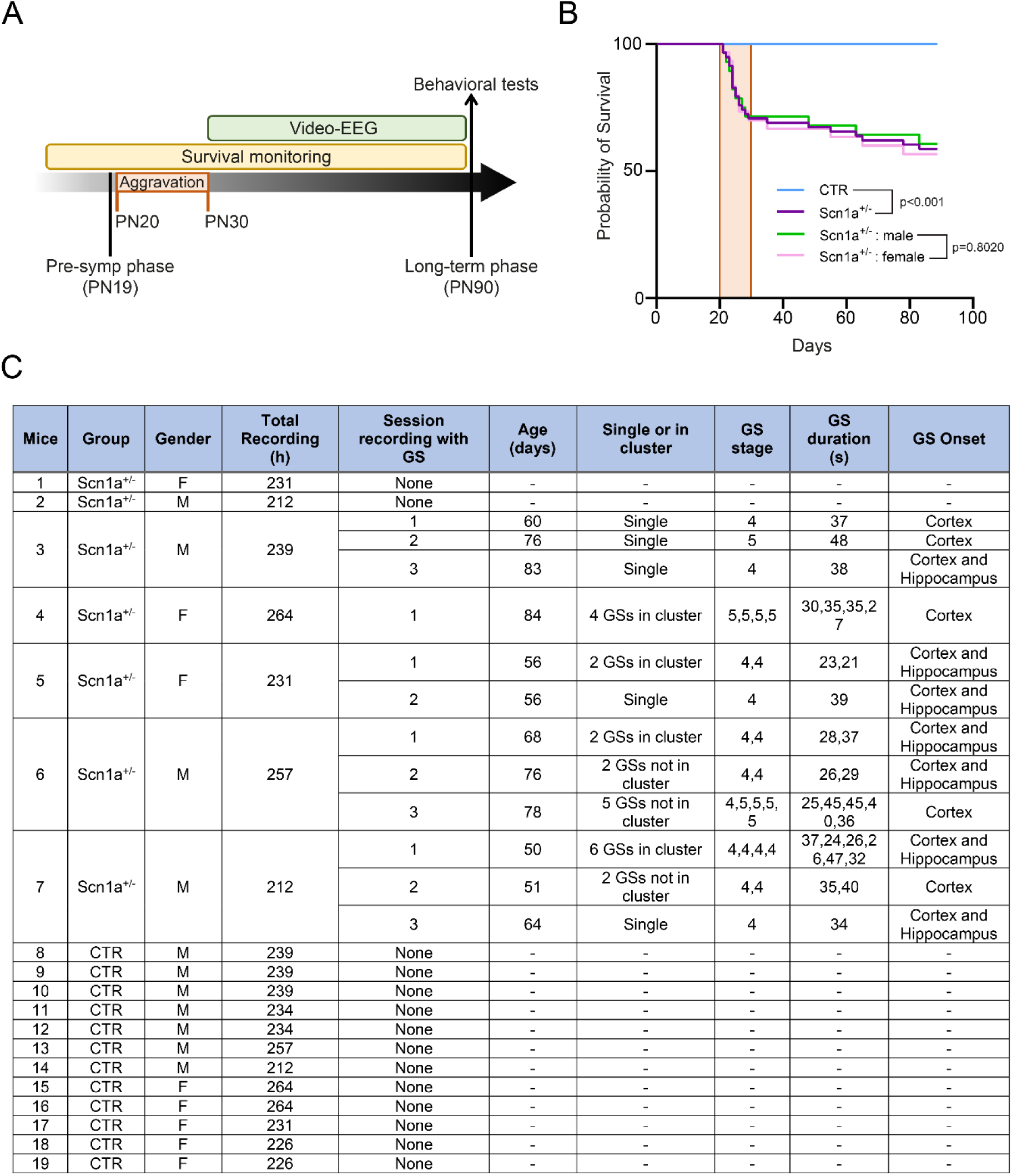
Kaplan-Meier and EEG details. **(A)** Schematic overview of the different phases of DS progression from the pre-symptomatic phase to the long-term phase, with the experimental timeline of results presented in Figure 1. **(B)** Kaplan-Meier survival curves showing the probability of survival (%) of the four groups of mice (CTR: n = 68; *Scn1a*^⁺/⁻^: n = 58, including males: n = 28 and females: n = 30). No significant difference was observed between male and female *Scn1a*^⁺/⁻^ mice (log-rank test, p = 0.8020), while survival was significantly reduced in *Scn1a*^⁺/⁻^ mice compared to control littermates (log-rank test, p < 0.001). A decrease in survival probability occurred during the worsening phase (light beige box from PN20 to PN30), reflecting the severity of seizures. **(C)** Characteristics of mice monitored by EEG. For each animal, the table reports the total recording time from PN50 to PN90, the number of recording sessions with generalized seizures (GS), and the age at each recording session. For each recording session, it indicates the number of seizures, whether they occurred in clusters, the seizure stage (according to the Racine scale), the duration, and the onset, which was either cortical or cortical and hippocampal together.

**Supplemental Figure 2.**
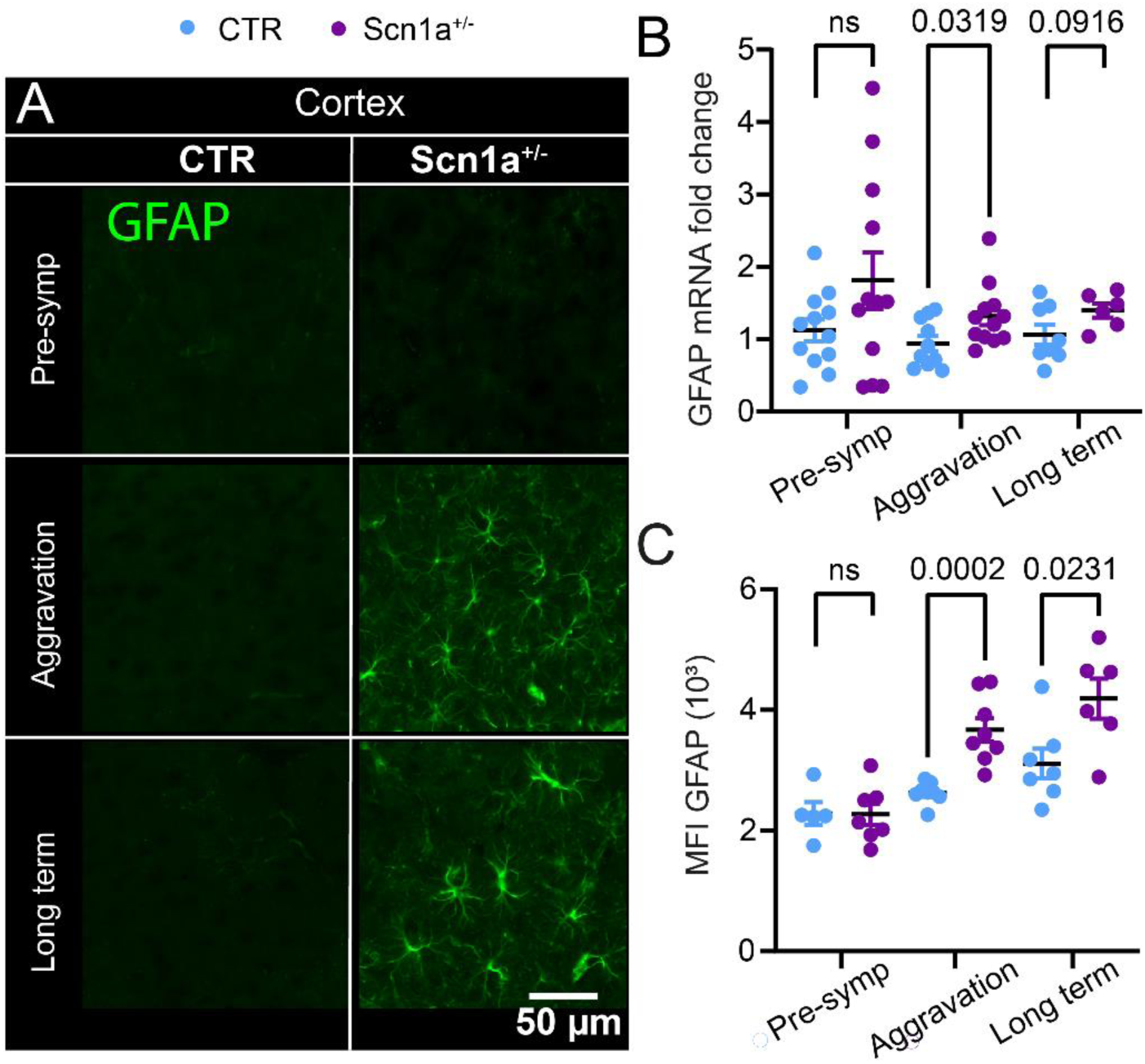
Long-term increases in cortical GFAP expression and astrocyte morphological remodeling in *Scn1a*^⁺/⁻^ mice. **(A)** Representative 40× GFAP images from the cortical area in CTR and *Scn1a*^⁺/⁻^ mice at pre-symptomatic, aggravation, and long-term phases. Scale bar: 50 µm. **(B–C)** Quantification of GFAP mRNA levels (B) and GFAP mean fluorescence intensity (MFI) (C) at pre-symptomatic, aggravation, and long-term phases. Unpaired t tests were performed at each time point: Pre-symptomatic: mRNA (CTR N=12; *Scn1a*^⁺/⁻^ N=12, p=0.1187, unpaired t test, t=1.624; df=22), MFI (CTR N=5; *Scn1a*^⁺/⁻^ N=7, p=0.9622), aggravation: mRNA (CTR N=10; *Scn1a*^⁺/⁻^ N=12, p=0.0319, unpaired t test, t=2.306; df=20), MFI (CTR N=8; *Scn1a*^⁺/⁻^ N=8, p=0.0002, unpaired t test, t=5.025; df=14). Long term: mRNA (CTR N=8; *Scn1a*^⁺/⁻^ N=6, p=0.0916, unpaired t test, t=1.833; df=12), MFI (CTR N=7; *Scn1a*^⁺/⁻^ N=6, unpaired t test, t=2.638; df=11).

**Supplemental Figure 3.**
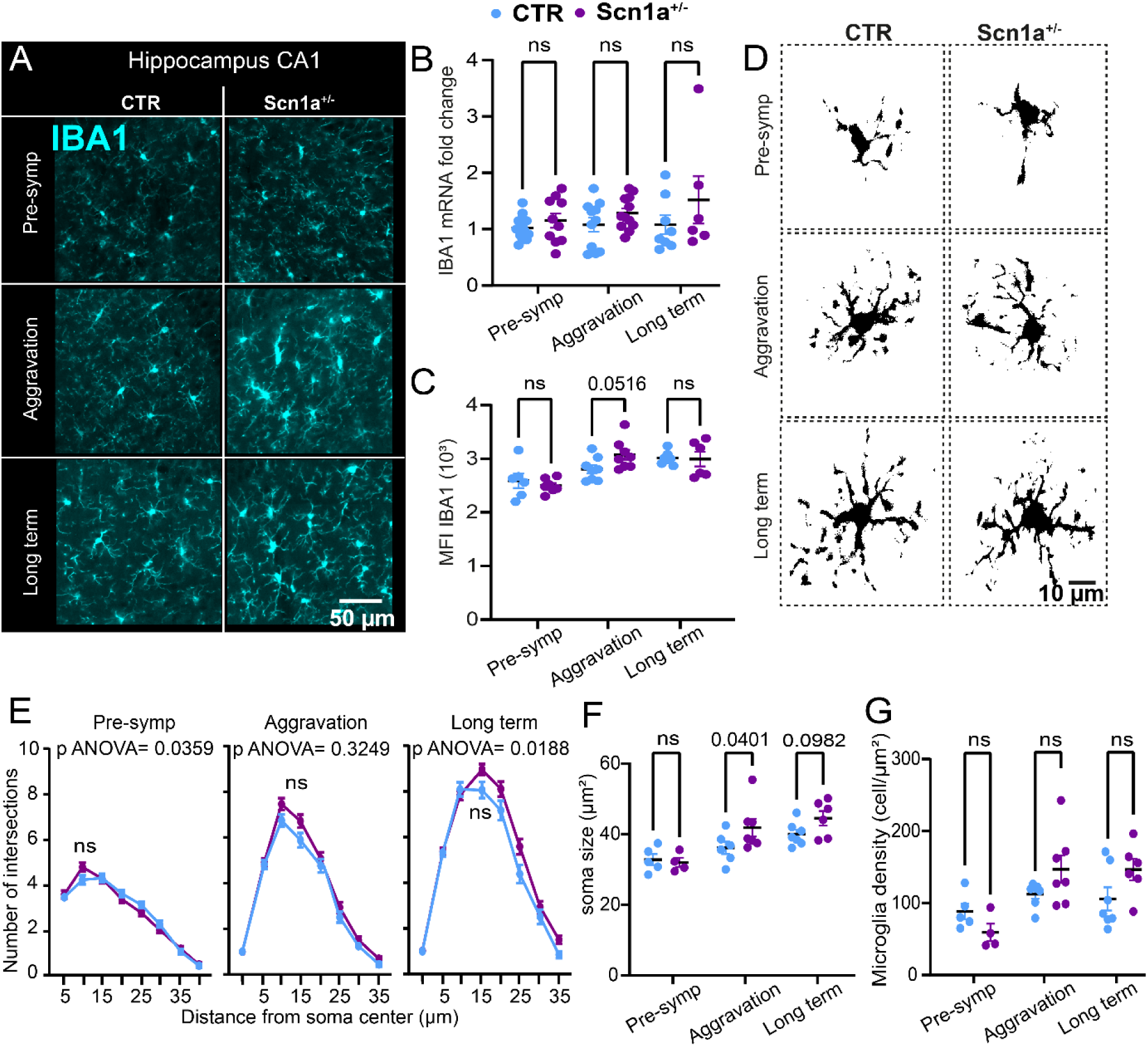
Patterns of hippocampal IBA1 expression, microglia morphology, and density in *Scn1a*^⁺/⁻^ mice. **(A)** Representative 40× IBA1 images from the dorsal hippocampal CA1 region in CTR and *Scn1a*^⁺/⁻^ mice. Scale bar: 50 µm. **(B–C)** Quantification of IBA1 mRNA levels (B) and IBA1 mean fluorescence intensity (MFI) (C) at pre-symptomatic, aggravation, and long-term phases. Mann-Whitney tests (mRNA) or Unpaired t tests (MFI) were performed at each time point: pre-symptomatic: mRNA (CTR N=12; *Scn1a*^⁺/⁻^ N=12, p=0.5492), MFI (CTR N=6; *Scn1a*^⁺/⁻^ N=7, p=0.5067; t=0.6864, df=11); Aggravation: mRNA (CTR N=11; *Scn1a*^⁺/⁻^ N=12, p=0.3236), MFI (CTR N=8; *Scn1a*^⁺/⁻^ N=8, p=0.0516; t=2.128; df=14); Long term: mRNA (CTR N=8; *Scn1a*^⁺/⁻^ N=6, p=0.2824), MFI (CTR N=7; *Scn1a*^⁺/⁻^ N=6, p=0.8965; t=0.1331, df=11). **(D–E)** Sholl analysis was performed on individual CA1 microglia at each stage. Intersections were quantified using concentric circles spaced 5 µm apart from the soma. Two-way ANOVA was used at each time point: pre-symptomatic: CTR 100 cells (N=5); *Scn1a*^⁺/⁻^ 100 cells (N=5); interaction p=0.0359, F(7, 1386)=2.152; followed by Bonferroni’s post hoc test; Aggravation: CTR 100 cells (N=5); *Scn1a*^⁺/⁻^ 100 cells (N=5); interaction p=0.3249, F(8, 1848)=1.152; Long term: CTR 100 cells (N=5); *Scn1a*^⁺/⁻^ 100 cells (N=5); interaction p=0.0188, F(8,1344)=2.304. followed by Bonferroni’s post hoc test. (F) Microglial soma size was quantified from IBA1 immunolabelling in the stratum radiatum. Mann-Whitney tests were performed at each time point: Pre-symptomatic: CTR N=5; *Scn1a*^⁺/⁻^ N=4; p=0.9048; Aggravation: CTR N=8; *Scn1a*^⁺/⁻^ N=7; p=0.0401; Long term: CTR N=5; *Scn1a*^⁺/⁻^ N=6; p=0.0982. (G) Microglial density quantified from IBA1 immunolabelling in the stratum radiatum. Mann-Whitney tests were performed at each time point: Pre-symptomatic: CTR N=5; *Scn1a*^⁺/⁻^ N=4; p=0.1111; Aggravation: CTR N=7; *Scn1a*^⁺/⁻^ N=7; p=0.1282; Long term: CTR N=5; *Scn1a*^⁺/⁻^ N=6; p=0.1375. For all datasets, each dot represents one mouse. Statistical significance: ***p < 0.001; ****p < 0.0001.

**Supplemental Figure 4.**
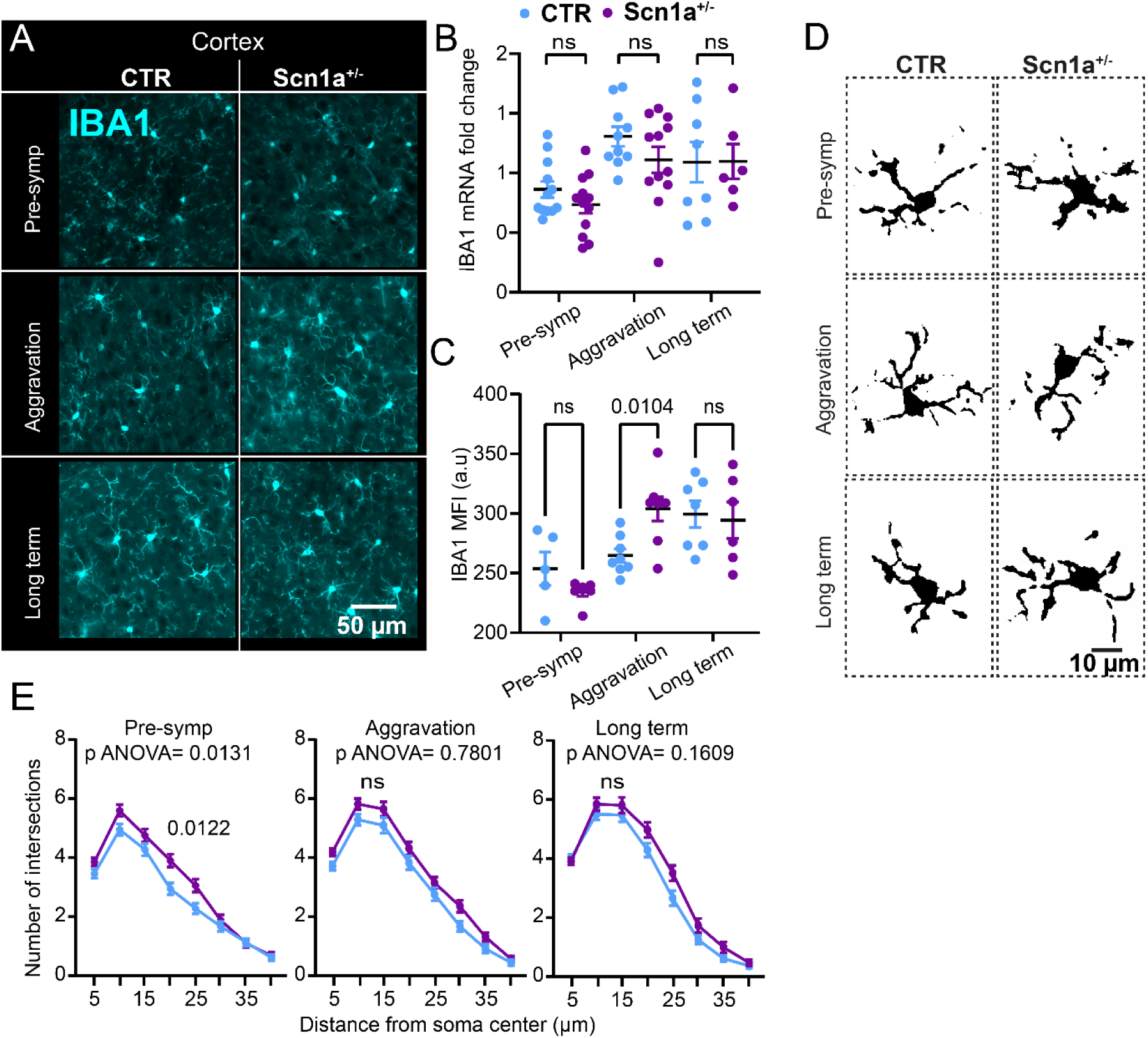
Patterns of cortical IBA1 expression, microglia morphology, and density in *Scn1a*^⁺/⁻^ mice. **(A)** Representative 40× IBA1 images from the cortex in CTR and *Scn1a*^⁺/⁻^ mice. Scale bar: 50 µm. **(B–C)** Quantification of IBA1 mRNA levels (B) and IBA1 mean fluorescence intensity (MFI) (C) at pre-symptomatic, aggravation, and long-term phases. **(B)** Unpaired t tests (mRNA) or Mann-Whitney tests (MFI) were performed at each time point: Pre-symptomatic: mRNA (CTR N=12; *Scn1a*^⁺/⁻^ N=12, p=0.2109; t=1.289, df=22), MFI (CTR N=6; *Scn1a*^⁺/⁻^ N=7, p=0.1881; Aggravation: mRNA (CTR N=11; *Scn1a*^⁺/⁻^ N=12, p=0.1801; t=1.389; df=20), MFI (CTR N=8; *Scn1a*^⁺/⁻^ N=8, p=0.0104; Long term: mRNA (CTR N=8; *Scn1a*^⁺/⁻^ N=6, p=0.9818; t=0.02327, df=12), MFI (CTR N=7; *Scn1a*^⁺/⁻^ N=6, p>0.9999; **(D–E)** Sholl analysis was performed on individual cortical microglia at each age. Intersections were quantified using concentric circles spaced 5 µm apart from the soma. Two-way ANOVA was used at each time point: Pre-symptomatic: CTR 100 cells (N=5); *Scn1a*^⁺/⁻^ 100 cells (N=5); interaction p=0.0131, F (5.240, 1038)=2.848; followed by Bonferroni’s post hoc test; Aggravation: CTR 100 cells (N=5); *Scn1a*^⁺/⁻^ 100 cells (N=5); interaction p=0.7801, F(4.517, 894.4)=0.4702; Long term: CTR 100 cells (N=5); *Scn1a*^⁺/⁻^ 100 cells (N=5); interaction p=0.1609, F(4.629, 823.9)=1.608.

**Supplemental Figure 5.**
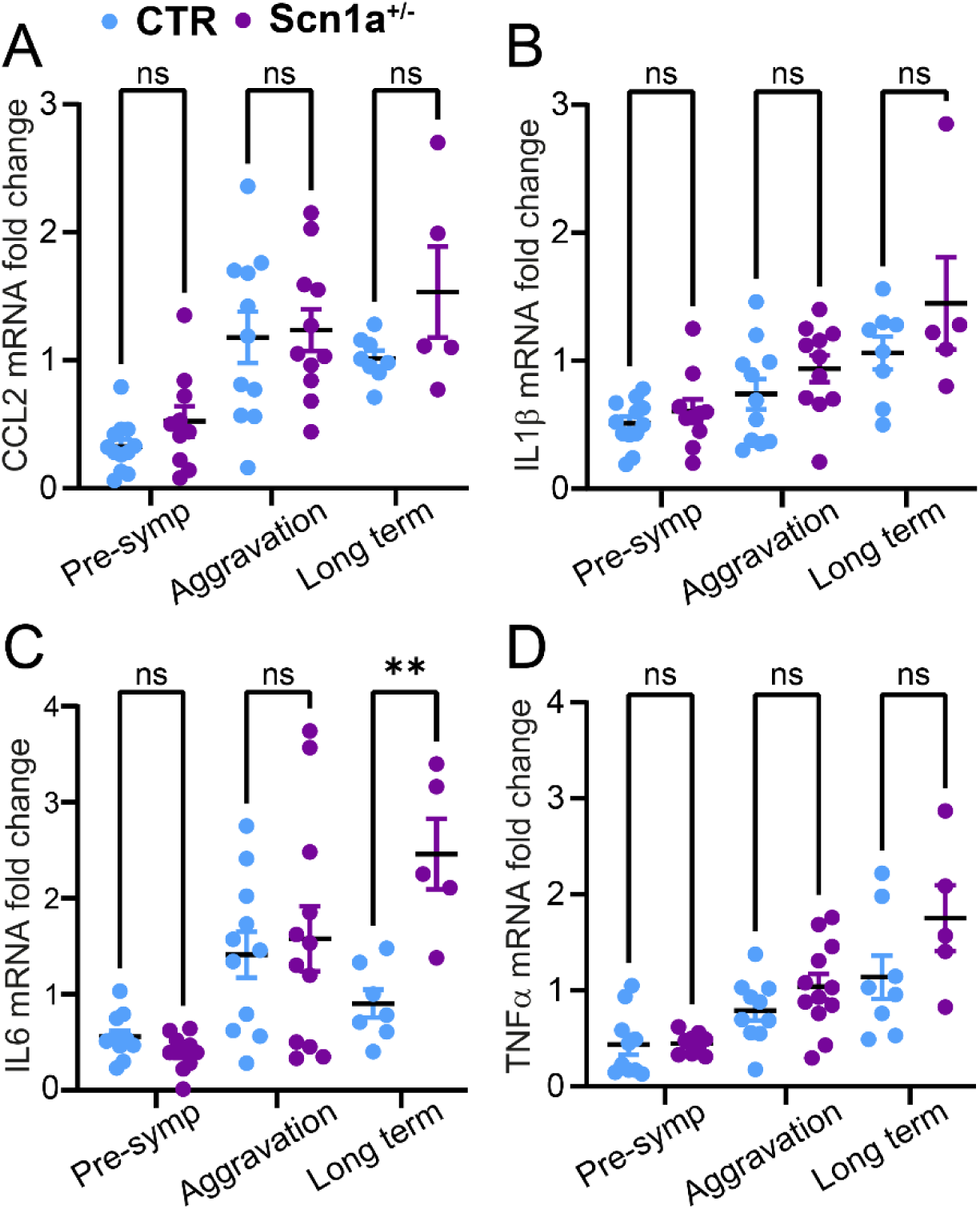
Profiles of pro-inflammatory cytokine expression during disease progression in the hippocampus of *Scn1a*^⁺/⁻^ mice. **(A-D)** Quantification of CCL2, IL1b, IL6, and TNFα mRNA expression levels. Mann–Whitney or t tests were performed at each time point: **(A)** CCL2: pre-symptomatic: CTR n=12; *Scn1a*^⁺/⁻^ n=10, p=0.1287, t=1.585; df=20. aggravation: CTR n=11; *Scn1a*^⁺/⁻^ n=11, p=0.8323, t=0.2145; df=20. Long term: CTR n=8; *Scn1a*^⁺/⁻^ n=5, p=0.0943, t=1.831; df=11. **(B)** IL1b: pre-symptomatic: CTR n=12; *Scn1a*^⁺/⁻^ n=10, p=0.5492. Aggravation: CTR n=11; *Scn1a*^⁺/⁻^ n=11, p=0.2169. Long term: CTR n=8; *Scn1a*^⁺/⁻^ n=5, p=0.6480. **(C)** IL6: pre-symptomatic: CTR n=12; *Scn1a*^⁺/⁻^ n=10, p=0.0747, t=1.881; df=20. aggravation: CTR n=11; *Scn1a*^⁺/⁻^ n=12, p=0.7007, t=0.3897; df=21. Long term: CTR n=7; *Scn1a*^⁺/⁻^ n=5, p=0.0013, t=4.422; df=10. **(D)** TNFα: pre-symptomatic: CTR n=12; *Scn1a*^⁺/⁻^ n=10, p=0.4585. aggravation: CTR n=11; *Scn1a*^⁺/⁻^ n=11, p=0.1854. Long term: CTR n=8; *Scn1a*^⁺/⁻^ n=5, p=0.1709.

**Supplemental Figure 6.**
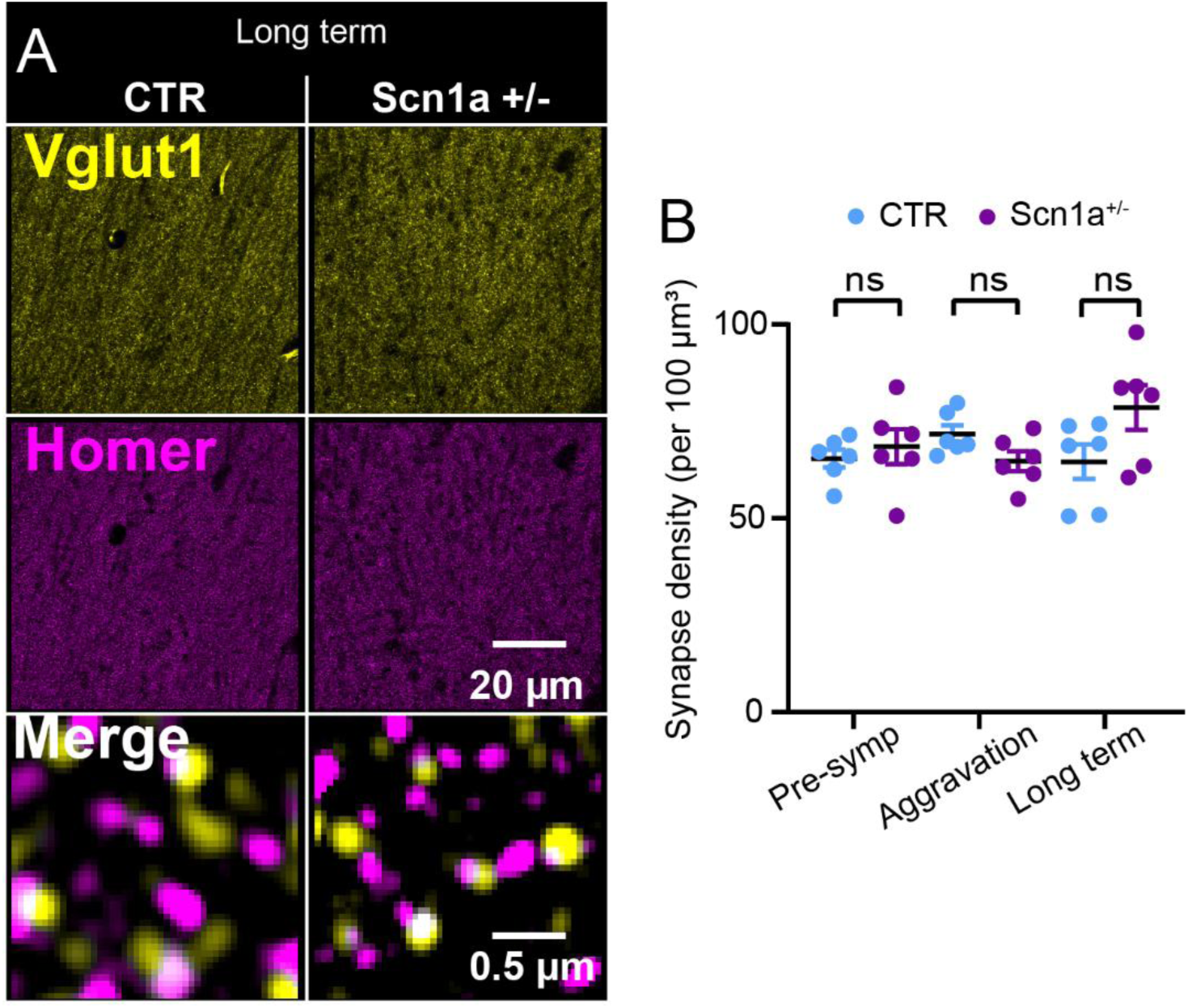
Synaptic density in CA1 is unchanged in *Scn1a*^⁺/⁻^ mice. **(A)** Representative 63× images of VGLUT1 and Homer puncta in the CA1 hippocampus of CTR and *Scn1a*^⁺/⁻^ mice. Scale bars: 20 µm (overview) and 0.5 µm (zoom). **(B)** Quantification of synaptic density at the pre-symptomatic, aggravation, and long-term phases. For all time points: CTR, N= 6; *Scn1a*^⁺/⁻^, N = 6. Mann–Whitney tests at each age: pre-symptomatic, p = 0.5887; aggravation, p = 0.0931; long term, p = 0.1320. Each dot represents a mouse.

**Supplemental Figure 7.**
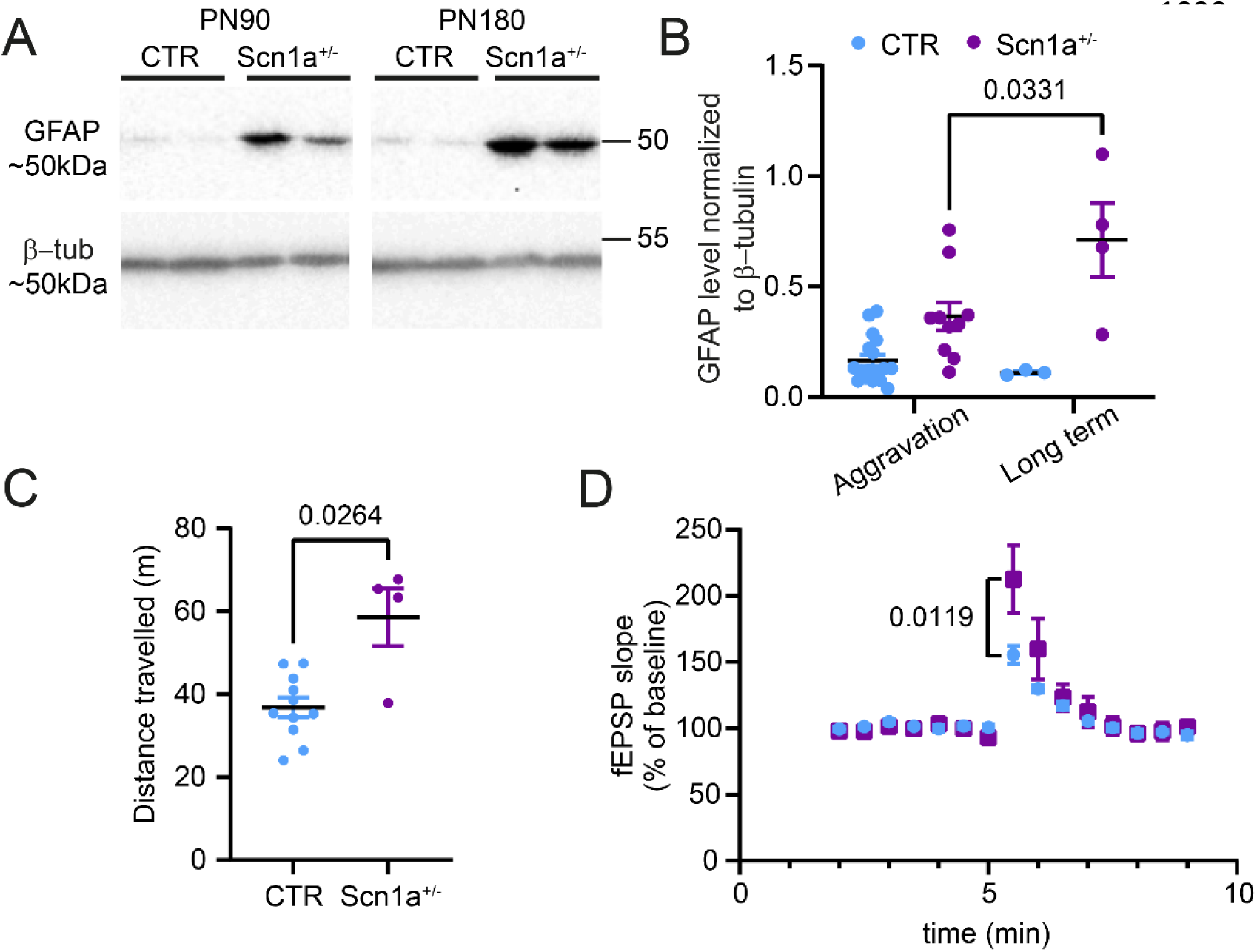
Increased GFAP expression and synaptic deficits are maintained >PN90 in *Scn1a*^⁺/⁻^ Mice. **(A-B)** Representative membrane and Western blot analysis of GFAP expression in the long-term phase and at 6 months old (PN180) of *Scn1a*^⁺/⁻^ Mice as compared to CTR. **(C)** Locomotion was assessed in the open field test at 6 months old (PN180). Mann-Whitney test, CTR N=11; *Scn1a*^⁺/⁻^ N=4; p=0.0264. **(D)** Post-tetanic potentiation (PTP) was increased in *Scn1a*^⁺/⁻^ mice (N = 3) compared to CTR (N = 6). Two-way ANOVA revealed a significant genotype × time interaction (p = 0.0383; F(1, 7) = 6.484). Bonferroni post hoc: p = 0.0119. Sample traces show averaged fEPSPs before and immediately after tetanization (dashed line).

**Supplemental Table 1.**
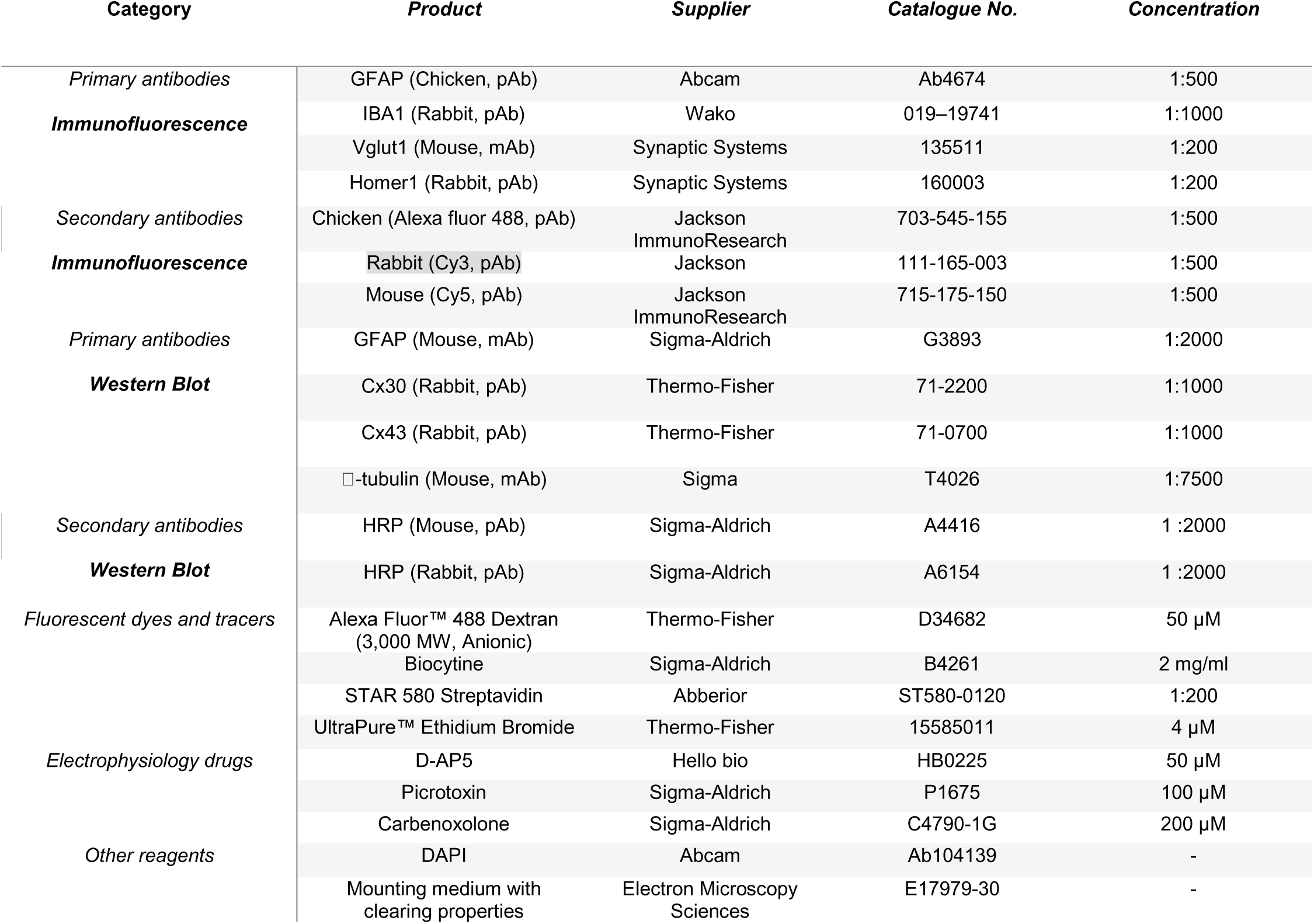
List of primary, secondary antibodies, and reagents.

**Supplemental Table 2.**
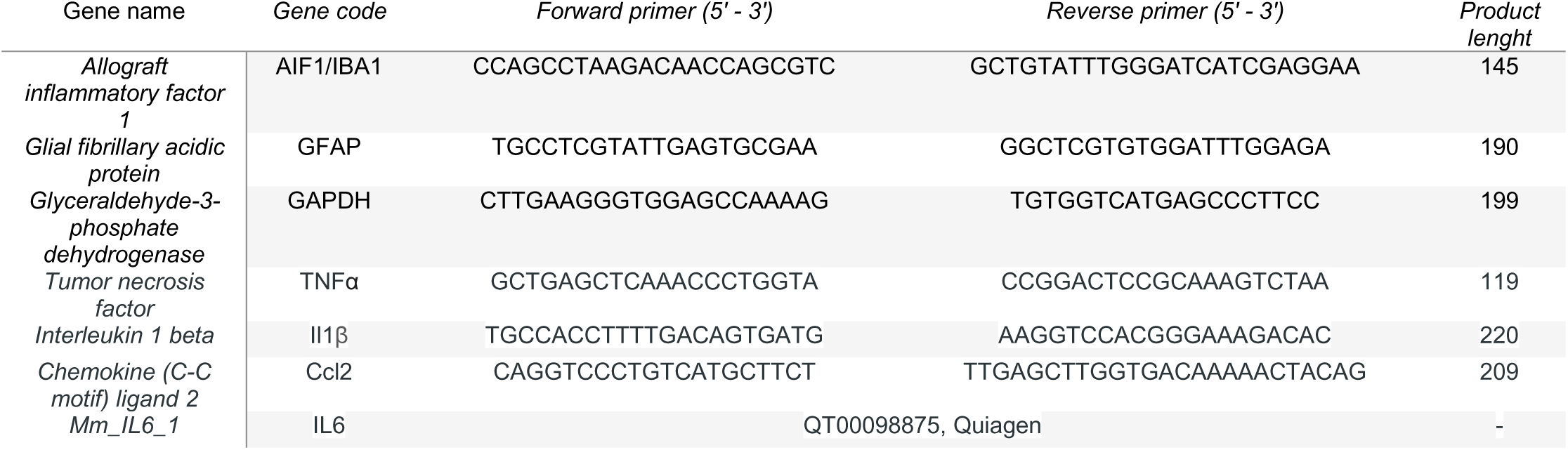
Primers used for qPCR.

