## Supplemental table 1 for "Long-lasting astrocyte remodeling in Dravet Syndrome *Scn1a*^+/–^ mouse model"

| Category | Product | Supplier | Catalogue No. | Concentration |
|---|---|---|---|---|
| Primary antibodies ***Immunofluorescence*** | GFAP (Chicken, pAb) | Abcam | Ab4674 | 1:500 |
| | IBA1 (Rabbit, pAb) | Wako | 019–19741 | 1:1000 |
| | Vglut1 (Mouse, mAb) | Synaptic Systems | 135511 | 1:200 |
| | Homer1 (Rabbit, pAb) | Synaptic Systems | 160003 | 1:200 |
| Secondary antibodies ***Immunofluorescence*** | Chicken (Alexa fluor 488, pAb) | Jackson ImmunoResearch | 703-545-155 | 1:500 |
| | Rabbit (Cy3, pAb) | Jackson | 111-165-003 | 1:500 |
| | Mouse (Cy5, pAb) | Jackson ImmunoResearch | 715-175-150 | 1:500 |
| Primary antibodies ***Western Blot*** | GFAP (Mouse, mAb) | Sigma-Aldrich | G3893 | 1:2000 |
| | Cx30 (Rabbit, pAb) | Thermo-Fisher | 71-2200 | 1:1000 |
| | Cx43 (Rabbit, pAb) | Thermo-Fisher | 71-0700 | 1:1000 |
| | β-tubulin (Mouse, mAb) | Sigma | T4026 | 1:7500 |
| Secondary antibodies ***Western Blot*** | HRP (Mouse, pAb) | Sigma-Aldrich | A4416 | 1 :2000 |
| | HRP (Rabbit, pAb) | Sigma-Aldrich | A6154 | 1 :2000 |
| Fluorescent dyes and tracers | Alexa Fluor™ 488 Dextran (3,000 MW, Anionic) | Thermo-Fisher | D34682 | 50 µM |
| | Biocytine | Sigma-Aldrich | B4261 | 2 mg/ml |
| | STAR 580 Streptavidin | Abberior | ST580-0120 | 1:200 |
| | UltraPure™ Ethidium Bromide | Thermo-Fisher | 15585011 | 4 µM |
| Electrophysiology drugs | D-AP5 | Hello bio | HB0225 | 50 µM |
| | Picrotixine | Sigma-Aldrich | P1675 | 100 µM |
| | Carbenoxolone | Sigma-Aldrich | C4790-1G | 200 µM |
| Other reagents | DAPI | Abcam | Ab104139 | - |
| | Mounting medium with clearing properties | Electron Microscopy Sciences | E17979-30 | - |

**Supplementary Table 1. Primers used for PCR amplification in this study**

| Protein | Antibody | Species/clonality | Company | Catalogue # /Lot# | Dilution (IHC, optimized) |
|---|---|---|---|---|---|
| **ERα** | **1D5** | Mouse/mAb | DAKO | M7047 / 00077836 | 1:150 |
| **ERβ** | **PPZ0506** | Mouse/mAb | Perseus Proteomics (Invitrogen, R&D) | 417100/A-2 | 1:600 |
| | **14C8** | Mouse/mAb | GeneTex | GTX70174/42142, 40784 | 1:1500 |
| | **PPG5/10** | Mouse/mAb | DAKO | M7292 / 00076093 | 1:60 |
| | | | BioRad | MCA1974G1 /161031 | |
| | | | Thermofisher | MA1-81281 / 74783060 | |
| | 6A12 | Mouse/mAb | Novus | NB200-303 / 1 | 1:225, 1:300 |
| | ab133467 | Rabbit/mAb | Abcam | ab133467 / GR97322-1 | 1:250 |
| | 68-4 | Rabbit/mAb | Upstate (Millipore) | 05-824 /n.d. | 1:300 |
| | ERb_503 | Chicken/pAb | *In-house produced* [2] | *n/a* | 1:900 |
| | H150 | Rabbit/pAb | Santa Cruz Biotechnology | sc-8974 / C1312 | 1:200 |
| | N-terminal | Rabbit/pAb | *In-house produced* [3] | *n/a* | 1:300 |
| | ab137381 | Rabbit/pAb | Abcam | ab137381 / GR106256-1 | 1:150 |
| | CT | Rabbit/pAb | Upstate (Millipore) | 07-359 /n.d. | 1:200, 1:400 |
| | HPA056644 | Rabbit/pAb | *In-house produced (HPA)* | *n/a* | 1:300 |
| | ab3577 | Rabbit/pAb | Abcam | ab3577 / GR73417-1 | 1:750 |
