## Supplemental table 2 for "Long-lasting astrocyte remodeling in Dravet Syndrome *Scn1a*^+/–^ mouse model"

| Gene name | Gene code | Forward primer (5' - 3') | Reverse primer (5' - 3') | Product lenght |
|---|---|---|---|---|
| *Allograft inflammatory factor 1* | AIF1/IBA1 | CCAGCCTAAGACAACCAGCGTC | GCTGTATTTGGGATCATCGAGGAA | 145 |
| *Glial fibrillary acidic protein* | GFAP | TGCCTCGTATTGAGTGCGAA | GGCTCGTGTGGATTTGGAGA | 190 |
| *Glyceraldehyde-3-phosphate dehydrogenase* | GAPDH | CTTGAAGGGTGGAGCCAAAAG | TGTGGTCATGAGCCCTTCC | 199 |
| *Tumor necrosis factor* | TNFα | GCTGAGCTCAAACCCTGGTA | CCGGACTCCGCAAAGTCTAA | 119 |
| *Interleukin 1 beta* | Il1β | TGCCACCTTTTGACAGTGATG | AAGGTCCACGGGAAAGACAC | 220 |
| *Chemokine (C-C motif) ligand 2* | Ccl2 | CAGGTCCCTGTCATGCTTCT | TTGAGCTTGGTGACAAAAACTACAG | 209 |
| *Mm_IL6_1* | IL6 | QT00098875, Quiagen | | - |

**Supplementary Table 1. Primers used for PCR amplification in this study**
