## Supplemental Extended Methods for "Long-lasting astrocyte remodeling in Dravet Syndrome *Scn1a*^+/–^ mouse model"

Supplemental Methods.

*Animal housing.*

We used the exon 25-floxed mouse line, which was maintained on a C57BL/6J background (Jackson Laboratories, Bar Harbor, ME). Animals were genotyped for the Scn1a floxed (F) allele using FHY311 (5’-CTTGATGTGTTGAAATTCAC-3') and FHY314 (5’-TATAGAGTGTTTAATCTCAAC-3'): WT allele, 846 BP; floxed allele, 1019 BP; and excised allele, 258 BP. Meox2-Cre mice (obtained from Jackson Laboratories) were received and maintained on a C57BL/6J background. Animals were genotyped for the presence of Cre using primers (5’-GGTTTCCCGCAGAACCTGAA-3') and (5’-CCATCGCTCGACCAGTTTAGT-3'). Mouse lines were independently maintained and mated together to generate F1. Both male and female mice were used for experiments. A batch of animals was examined at 6 months old for biochemistry, behavior, and electrophysiology experiments.

*In vivo EEG recordings and analysis.*

A bipolar electrode consisting of two isolated and twisted tungsten wires was inserted into the hippocampus (AP -2.0 mm, ML +1.5 mm, DV -2 mm, according to Paxinos and Franklin). For cortical recordings, two micro-screws (Pinnacle Technology Inc, #8209) were anchored in the same hemisphere, one in the frontal cortex (AP +1.7 mm, ML –1.0 mm, DV –1 mm) and the other in the parietal cortex (AP -2 mm, ML –2.0 mm, DV –1 mm) for fronto-parietal recordings. All wires were connected to a prefabricated micro-connector (6-Pin Surface Mount Connector for Mouse, Pinnacle Technology Inc, #8235-SM), secured and insulated using dental acrylic cement (Palladur). Video/EEG recordings were performed using Sirenia acquisition software (Pinnacle Technology Inc, Lawrence, KS, USA) with an analog low-pass filter (250 Hz).

One week after surgery, the mice were habituated to circular Plexiglas EEG recording chambers (Rohm, Philadelphia, Pennsylvania, USA), followed by a second session connected to the EEG preamplifier to acclimate them to its use. Recordings were then made continuously every two days, alternating between night and day, from PN50 to approximately PN90. This program optimized animal welfare over time and minimized implant loss, particularly given the long recording duration. Each recording session was performed at fixed times (from 8 am to 8 pm and from 8 pm to 8 am). For EEG analyses and quantification, EEG traces were analyzed together with synchronized video recordings in a blinded manner. Seizure events were first automatically detected using Neuroscore, excluding time segments previously identified as containing artifacts. These artifacts (including signal instability caused by hardware issues, head-mount scratching, or electrical interference from a water bottle) were visually identified and manually removed. Detected seizure events were then qualitatively reviewed to determine concordance with canonical EEG seizure features, such as seizure onset marked by pre-ictal spikes and seizure termination associated with EEG flattening. Final validation of seizure events was performed systematically by cross-referencing EEG findings with modified Racine scale–associated behaviors observed on synchronized video recordings. Racine scale used, 0: No behavioral response; 1: motionless staring (with orofacial automatism); 2: Head nodding; 3: Unilateral forelimb clonus; 4: Bilateral forelimb myoclonus with rearing; 5: Tonic-clonic with rearing and falling. Details of the seizures (recording times, clustering, duration, stage, and seizure onset) are reported in Supplementary Fig.1C.

*Ex vivo extracellular field recordings*

The slices were perfused with aCSF containing picrotoxin (100 μM) at 30–32 ◦C at a flow rate of 3 ml/min. A cut between CA3 and CA1 was made to avoid epileptiform activity. EPSPs were evoked with a concentric bipolar electrode (FHC, SKU 30200) and recorded with an aCSF-filled glass pipette (1-2 MΩ), both placed in the stratum radiatum of CA1 at least 200 μm apart from each other. Extracellular field EPSPs (fEPSPs) were recorded and filtered (low-pass at 1 kHz) with an Axopatch 200B or a MultiClamp 700B amplifier (Molecular Devices, San Jose, CA, USA), digitized at 10 kHz with an A/D converter (Digidata 1322 A, Axon Instruments), then stored and analyzed on a computer using Pclamp11 software (Molecular Devices, San Jose, CA, USA). Input-output (I–O) relationships for fEPSPs were measured at the start of each experiment by applying a series of stimuli of increasing intensity to Schaffer's collaterals and plotting the initial slope of the fEPSP against the fiber-volley amplitude. Paired-pulse facilitation (PPF) was evoked by administering two stimuli at a 40 ms interval and was measured by dividing the maximum amplitude of the second response by that of the first. Long-term potentiation (LTP) was induced by tetanic stimulation of Schaffer collaterals (two trains of 100 Hz for 1 s, 20 s apart). Post-tetanic potentiation (PTP) was induced with the same tetanic stimulation, but in the presence of D-(-)-2-Amino-5-phosphonopentanoic acid (D-AP5, 50 μM, NMDA receptor antagonist, Hello Bio HB0225). The readily releasable vesicle pool was evaluated using a 10 Hz, 30 s stimulation protocol and normalized to baseline (5 min).

*Whole-cell patch clamp recording, dye loading, and image acquisition*

The identity of the patched cells was confirmed by applying positive and negative voltage steps (from -80 to +40 mV), which consistently showed a linear current-voltage relationship with a low slope resistance (10-15 MΩ) and no signs of voltage-activated currents, as expected for astrocytes. A gap junction-impermeant dye was used to eliminate the possibility that observed fluorescent intensity changes were due to diffusion of dye into other astrocytes through gap junctions. Astrocyte whole-cell membrane voltages and currents were recorded and filtered (1kHz) with an Axopatch 200B amplifier (Molecular Devices, Foster City, CA), sampled at 5kHz with an A/D converter (Digidata 1322A; Molecular Devices, Foster City, CA), and stored and analyzed on a computer with Pclamp11 software (Molecular Devices, Foster City, CA).

To assess the 3D morphology of astrocytes, recorded cells were held at −80 mV and filled with the gap-junction-impermeant dye Alexa Fluor 488 for 10 min. Then, the slices were fixed at 4°C for 2h in 4% paraformaldehyde in PBS (Ready-to-Use Fixative Solution, Biotium 22023), pH 7.4, and washed three times in 10 mM PBS for 10 minutes each before being mounted on glass slides using a clearing mounting medium. Image stacks were acquired with 488 nm laser-line excitation using a SP8 confocal microscope (Leica) in resonant-scanning mode at 8 kHz (40X magnification; 1024 x 1024 with 32-line average). 3D astrocytes morphological reconstruction: Astrocytes were reconstructed in three dimensions using IMARIS software (version 10.9, Bitplane). Confocal image stacks were imported into IMARIS, and individual astrocytes were segmented using the software's filament-tracing and surface-rendering tools. Reconstructions allowed visualization and quantitative analysis of astrocytic morphology, branching, territorial volume, and soma volume. To evaluate coupling, astrocytes were loaded passively with biocytin for 10 min in current-clamp mode. The slices were then fixed at 4°C for 12 h in 4% paraformaldehyde in PBS (Ready-to-Use Fixative Solution, Biotium 22023, pH 7.4), then stored in PBS. Slices were washed three times in 10 mM PBS solution for 10 minutes each. Streptavidin conjugated to STAR at 1:200 dilution in solution with 0.1% Triton and 0.2% Gelatin was used to bind biocytin for 2 hours. Slices were then washed three times in PBS solution for 10 minutes each. The slices were mounted on glass slides with ProLong Gold Antifade Mountant (Thermo Fisher Scientific, Waltham, MA, USA), and clear nail polish was applied around the perimeter of the coverslip. Image stacks were acquired with 580 nm laser-line excitation using a SP8 confocal microscope (Leica), at 40× magnification and 1024×1024 resolution, centered on the patched soma. Acquisition covered the full depth of the slice, from the surface down to where the fluorescence signal was no longer detectable, and extended laterally across the hippocampal region containing the labeled astrocytic network. 3D coordinates of each cell were extracted from the resulting image stacks. For each sample, the x-, y-, and z-positions of astrocyte somata were recorded in micrometers. Data processing and convex hull analysis were performed using the Python pandas library. The spatial distribution of astrocytes was analyzed by computing the Convex Hull using the scipy spatial ConvexHull function, with the QJ option to account for near-coplanar points. The Convex Hull volume and surface area were used to estimate the spatial extent of each astrocytic network. Distance and connectivity analysis: Pairwise Euclidean distances between astrocyte somata were calculated using the scipy.spatial.distance.pdist and squareform functions. Each astrocyte was modeled as a sphere with a 40 or 50 µm diameter for the aggravation and the long-term phases, respectively. Two astrocytes were considered connected if the distance between their centers was less than the sphere diameter. From this connectivity matrix, the number of neighbors for each astrocyte and the mean distance to connected neighbors were computed. Visualization: 3D scatter plots of astrocyte positions were generated using matplotlib, with each point colored according to its number of neighbors. Convex Hull boundaries were superimposed to provide a visual reference of the network volume.

*qPCR and quantification.*

Brain tissues from mice were kept dry frozen until the extraction day. Tissues were lysed within Lysing Matrix D tube in the Fast-Prep sample preparation system (MP Biomedicals, Santa Ana, USA 6913500). Tissue lysates were homogenised on QIAshredder columns (Qiagen 79656). Total RNA from tissues was extracted and purified using the RNeasy Plus Mini Kit (Qiagen 74136). The RNA concentration of each sample was measured with a Nanodrop 2000c spectrophotometer (Thermo Fisher Scientific, Waltham, USA). For each sample, 1 µg of total RNA was reverse-transcribed using random hexamers of the Transcriptor Universal cDNA Master (Roche, Pleasanton, USA, 05893151001). RT-qPCR was conducted using 2 µL cDNA (10 ng), 500 nmol/L of forward and reverse primers, and 2.5 µL of SYBR Green PCR Master Mix (Roche 04887352001) in triplicate wells. Amplification was performed on a LightCycler 480 apparatus (Roche) with 45 cycles (95°C/10 sec, 60°C/10 sec, 72°C for 1 sec). After amplification, the melting curve was assessed to ensure the presence of a single product. Each plate carried serial dilutions of an RNA calibrator to generate a standard curve of the primer's efficiency during the run. The levels of cDNA were quantified by the comparative 2ΔΔCt method. Ct values of the target gene were normalized to the Ct values of the housekeeping gene GAPDH (glyceraldehyde-3-phosphate dehydrogenase). Each target value is expressed as an n-fold difference relative to the control group. Primers are listed in Supplemental Table 2.

*Behavioral tests.*

Mice were housed in group cages with 4–3 individuals per cage, supplemented with enrichment cotton nests and nest boxes. Behavioral experiments were performed between 8:00 am – 4:00 pm in a light color wall room with control light devices, with the experimenter out of the animal's sight. Behavioral experiments were performed on animals between PN85 and PN95. Marble burying: mice were placed individually in a clean cage with 10 cm of bedding and 15 marbles evenly distributed on top. After 30 minutes in the cage, the mice were removed. We evaluated marble burial using the following Score: 0 for a buried marble, 1 for a half-buried marble, and 2 for a visible marble. Open field: mice were tested in the open field apparatus (50 cm × 45 cm). The distance travelled was recorded using the EthoVision XT video-tracking system (Noldus, Wageningen, Netherlands). Each mouse was placed at the periphery of the open field and allowed to freely explore the apparatus for 10 min, with the experimenter out of the animal's sight. Three Chambers Social preference test: The apparatus is a transparent cage with a central starting compartment and two side compartments, each with a circular grid cup (goal box) at its extremity, where the congener and the object can be placed during testing. The positions of the congener and object boxes were counterbalanced to avoid potential spatial preference. First, the mouse was placed in the central box and allowed to freely explore the apparatus for 10 min. Two minutes later, a C57Bl/6J juvenile congener from the same sex was placed in one goal box, and an object was placed in the opposite compartment. The mouse was then placed in the starting central compartment and allowed to explore the apparatus freely for 10 min. The duration in chambers was automatically measured. Y-maze test: The apparatus used to test spatial working memory was a Y-maze made of plexiglass with three identical arms (40×9×16 cm) arranged at 120° to each other. Each arm had walls with specific motifs that allowed it to be distinguished from the others. Each mouse was placed at the end of one of the three arms and allowed to explore freely the apparatus for 8 minutes. Alternations were automatically measured as successive entries into each of the three arms within overlapping triplet sets (e.g., ABC, BCA …) and calculated as an index of working memory performance.

*Immunofluorescence and quantifications.*

For microglial soma size quantification, a threshold was applied to select only microglial somata (IJ_IsoData, lower threshold = 700; upper threshold = maximum intensity). Particle analysis was then performed in FIJI by using a size constraint between 10 μm² and infinity. For each particle detected, the cell body surface area was measured automatically. A total number of 2,252 microglia for CTR and 1,505 microglia for Scn1a$^{+/-}$ from 4 and 6 mice, respectively for the pre-symptomatic phase, 5386 microglia for CTR and 6340 microglia for Scn1a$^{+/-}$ from 5 and 6 mice respectively for the aggravation phase, and 4225 microglia for CTR and 6295 microglia for Scn1a$^{+/-}$ from 5 and 6 mice respectively for the long-term stabilization phase. For synapses quantification: Antibodies against vesicular transporter 1 (Vglut1) and Homer protein homolog 1 (Homer1) were used to label pre- and post-synapses, respectively. The secondary antibodies used were anti-rabbit IgG conjugated to Cy3 and anti-mouse conjugated to Cy5. For synapse analyses, images were acquired with a confocal microscope (Zeiss Airyscan, Plateforme Montpellier Ressources Imagerie) using a 63X oil immersion objective (image size: 103.10× 103.10 μm, Resolution x,y: 140 nm; z: 450 nm). Two regions of interest in two sections were acquired per animal. For each section, in both channels (488 nm and 633 nm), we imaged serial optical sections at 0.17 μm intervals over a total of 5 μm, for a total of 30 optical sections. For image analysis, automated quantification was performed using the Distance Analysis plugin

(DiAna) (REF). The plugin performs automated object-based co-localization and distance analysis in 3D. As the first step, we used the global intensity thresholding (median) to select dots in each channel corresponding to Vglut1 and Homer1. This step created a new image for each channel representing each dot. Then, the plugin evaluates the co-localization between the dots of each channel. Co-localized signals were considered synapses, and the number of synapses was automatically calculated.

*Western blots.*

Hippocampi were homogenized for 30 min in 1.5 mL Eppendorf tubes using a plastic pestle (Argos Technologies, 7339-901) in Mesoscale lysis buffer (R60TX-3) supplemented with protease and phosphatase inhibitors (R70AA-1). Homogenates were then centrifuged at 16,000 × g for 15 min at 4°C, and the resulting supernatants were immediately stored at −80°C. Proteins were resolved on 12% SDS-polyacrylamide gels. The resolving gel was prepared using distilled water, 1.5 M Tris–HCl (pH 8.8, Biorad #1610771), 10% SDS (Euromedex #EV0660), 40% acrylamide/bis-acrylamide (Biorad #1610148), 10% APS (Ammonium perfulfate), and TEMED (sigma Aldrich #T7024), and allowed to polymerize for ~45 min. A 4% stacking gel (0.5 M Tris–HCl, pH 6.8) was then poured and polymerized for ~30 min before electrophoresis. Following gel polymerization, electrophoresis was performed in running buffer. Molecular weight markers and samples were loaded (3 µL of the PageRuler ladder per lane, PageRuler, PI26619, Thermo Scientific). Proteins were separated for approximately 1h 20 min at 0.12 A. Transfers on membranes (nitrocellulose, Amersham protran #10600019) were carried out at 100 V for ~1 h in a transfer buffer containing 30% ethanol. Following transfer, membranes were cut according to the molecular weights of the proteins of interest. Membranes were blocked for 1 h at room temperature in PBS-T containing 5% non-fat milk, then washed three times for 5 min in PBS-T. Membranes were incubated with primary antibodies diluted in SuperBlock (Thermo Scientific #37515) containing 0.01% sodium azide for 1 h at room temperature, followed by overnight incubation at 4 °C. Antibodies against Cx30, Cx43, GFAP, and β-tubulin were employed. After washing (3 × 5 min in PBS-T), membranes were incubated with horse radish peroxidase (HRP)- conjugated secondary antibodies diluted in PBS-T with 5% milk for 2h at room temperature. Membranes were then rewashed (3 × 5 min, PBS-T) prior to detection. Protein signals were detected using ECL (ECL Crescendo Millipore #WBLUR0500) substrate on a Bio-Rad ChemiDoc system, and images were saved for analysis (FIJI). If required, membranes were stripped by incubating in 5 mL ReBlot (Millipore #2504) solution diluted in 45 mL distilled water for 30 min at room temperature with agitation, followed by re-blocking and re-probing as described above.
