## Supplementary figures and images for "Long-lasting astrocyte remodeling in Dravet Syndrome *Scn1a*^+/–^ mouse model"

### Supplemental figure 1

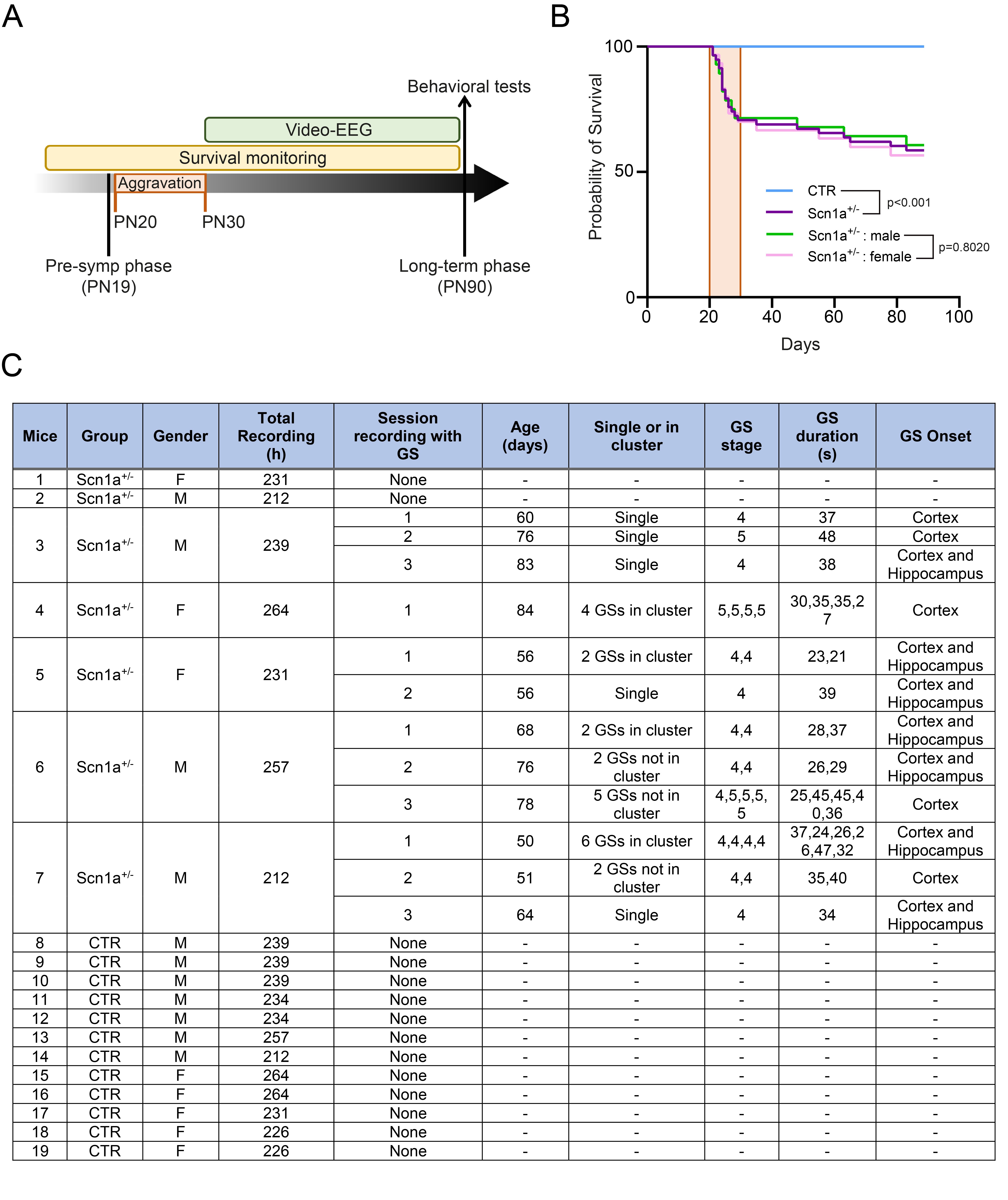

### Supplemental figure 2

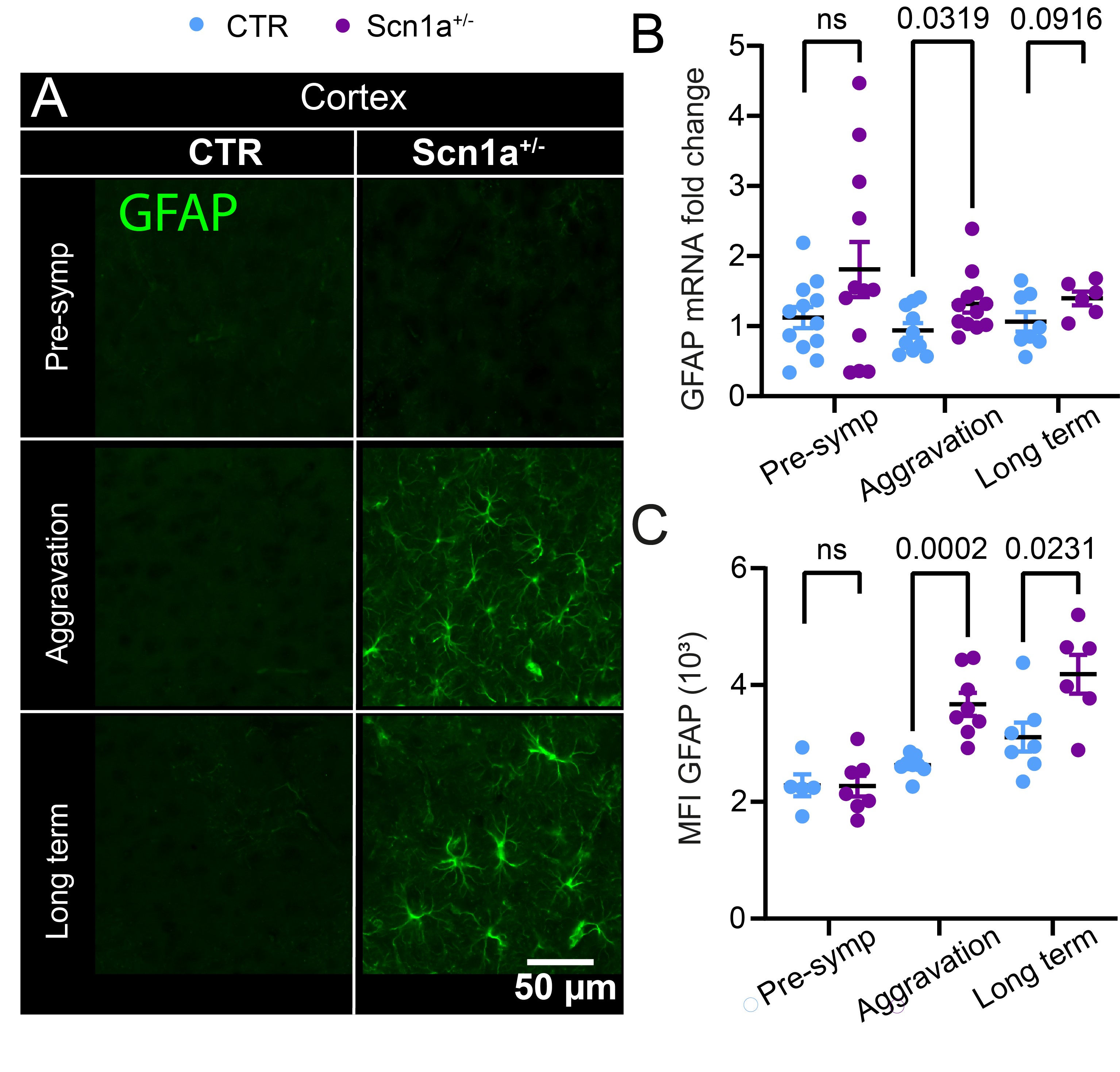

### Supplemental figure 3

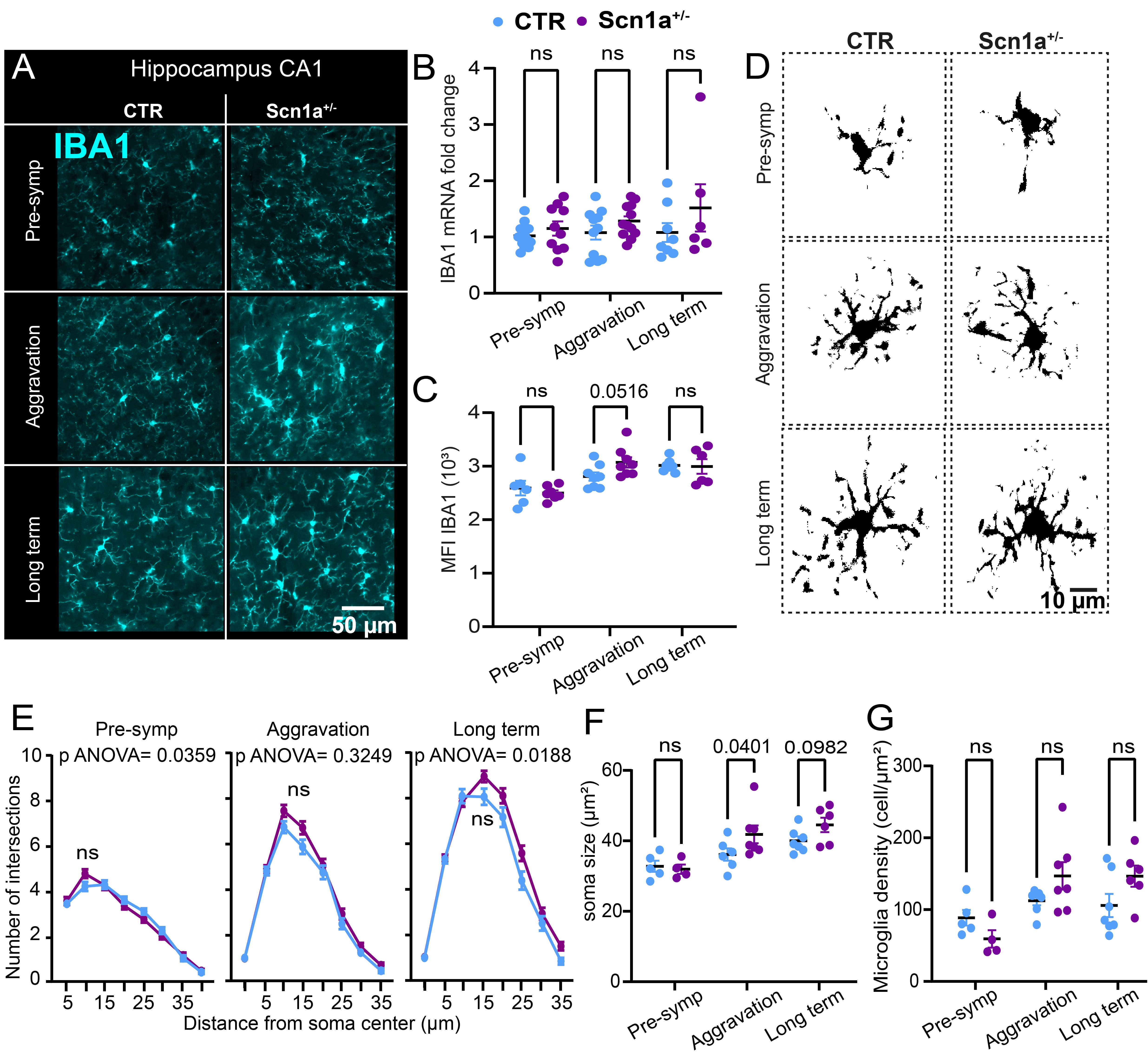

### Supplemental figure 4

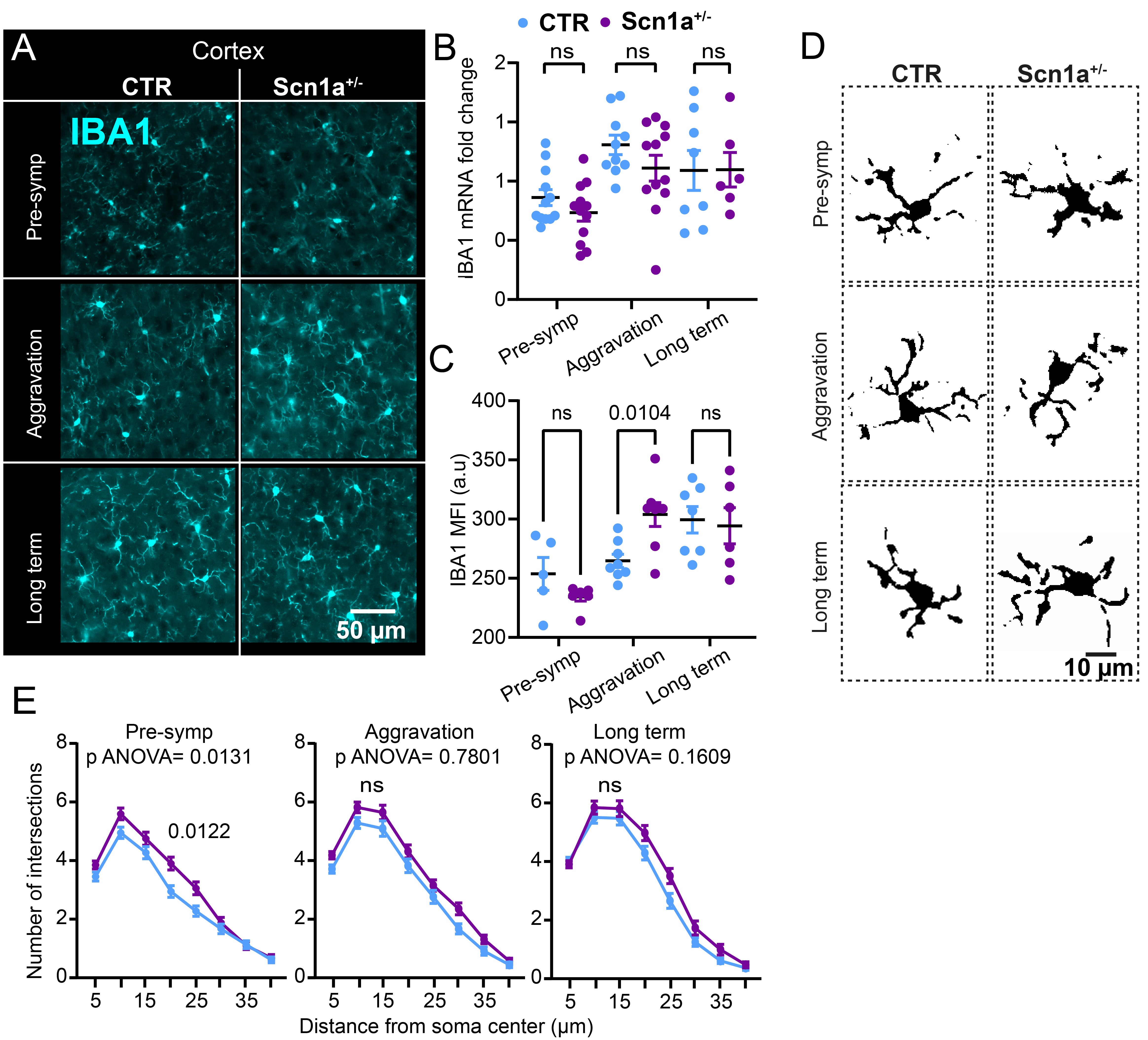

### Supplemental figure 5

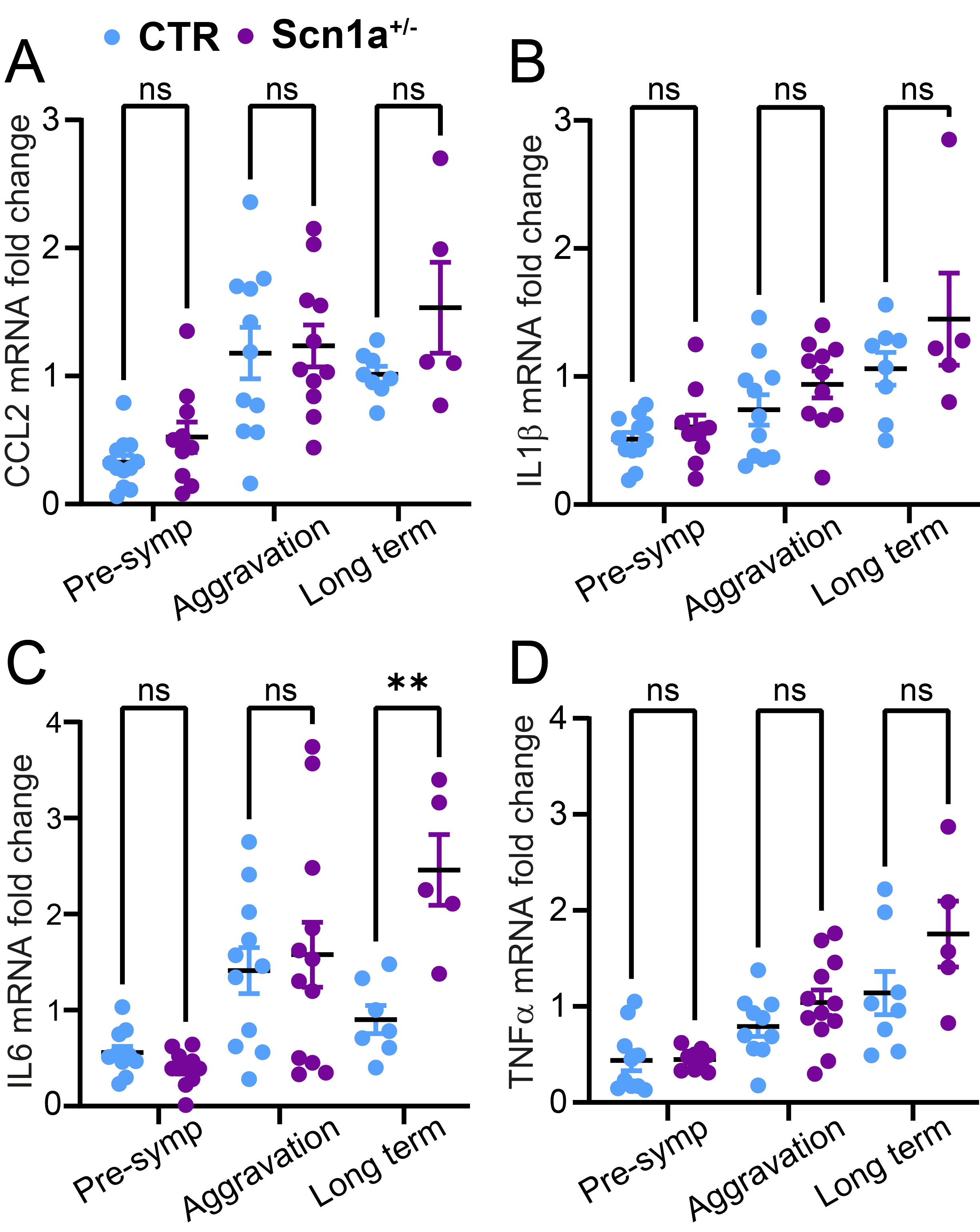

### Supplemental figure 6

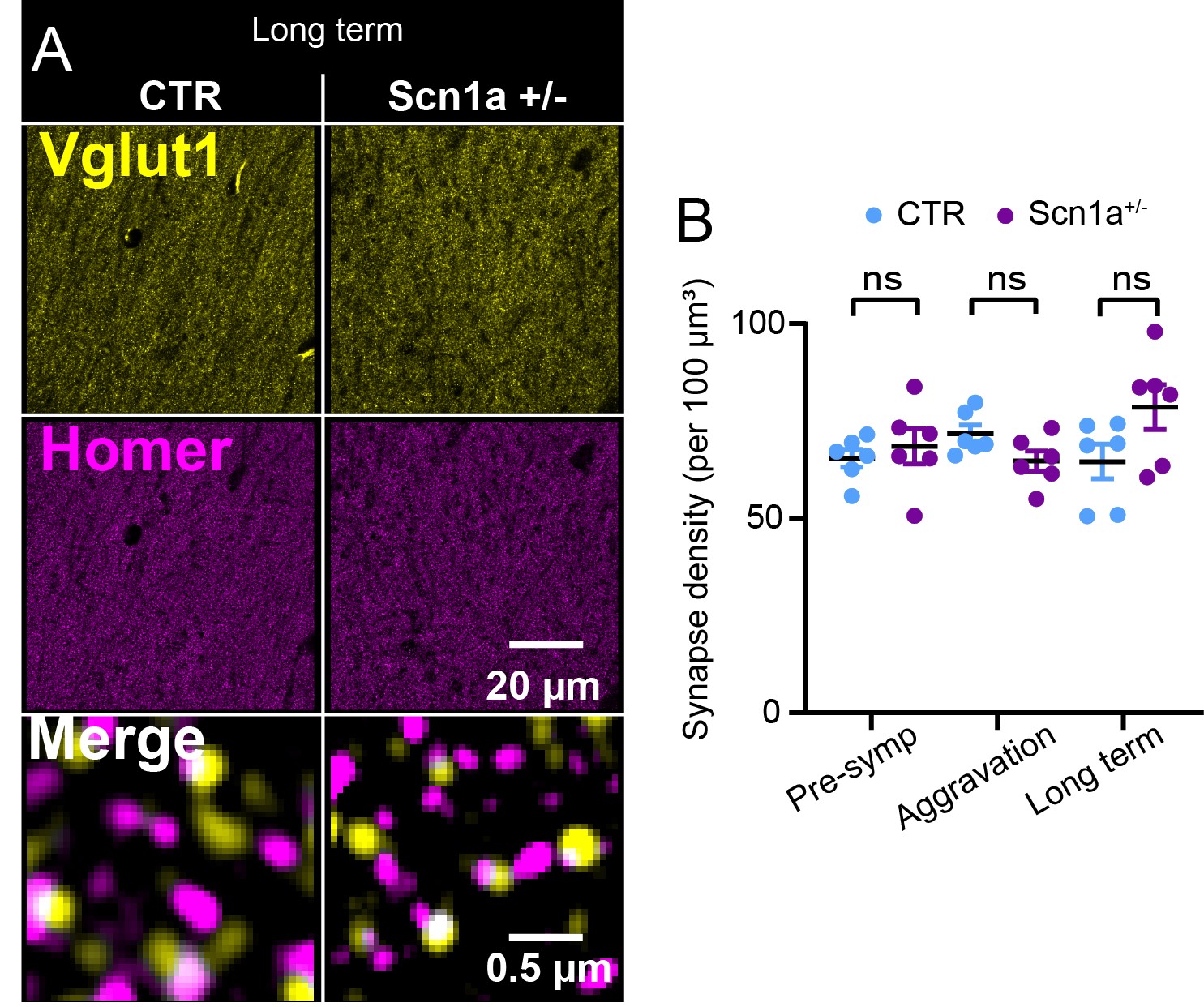

### Supplemental figure 7

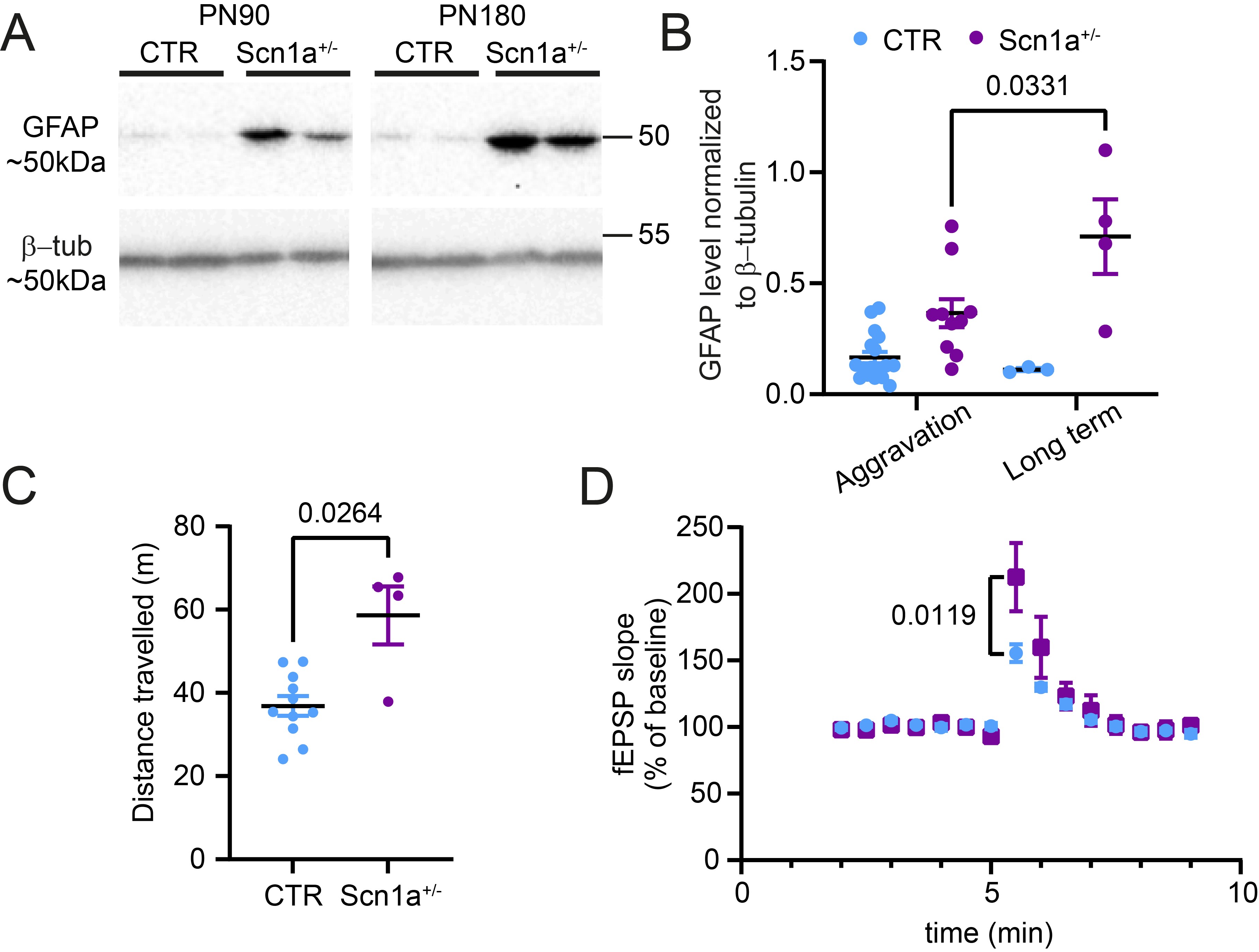
